# Microbiota produced indole metabolites disrupt host cell mitochondrial energy production and inhibit *Cryptosporidium parvum* growth

**DOI:** 10.1101/2023.05.25.542157

**Authors:** Lisa J. Funkhouser-Jones, Rui Xu, Georgia Wilke, Yong Fu, Lawrence A. Shriefer, Heyde Makimaa, Rachel Rodgers, Elizabeth A. Kennedy, Kelli L. VanDussen, Thaddeus S. Stappenbeck, Megan T. Baldridge, L. David Sibley

## Abstract

Cryptosporidiosis is a leading cause of life-threatening diarrhea in young children in resource-poor settings. Susceptibility rapidly declines with age, associated with changes in the microbiota. To explore microbial influences on susceptibility, we screened 85 microbiota- associated metabolites enriched in the adult gut for their effects on *C. parvum* growth in vitro. We identified eight inhibitory metabolites in three main classes: secondary bile salts/acids, a vitamin B_6_ precursor, and indoles. Growth restriction of *C. parvum* by indoles did not depend on the host aryl hydrocarbon receptor (AhR) pathway. Instead, treatment impaired host mitochondrial function and reduced total cellular ATP, as well as directly reduced the membrane potential in the parasite mitosome, a degenerate mitochondria. Oral administration of indoles, or reconstitution of the gut microbiota with indole producing bacteria, delayed life cycle progression of the parasite in vitro and reduced severity of *C. parvum* infection in mice. Collectively, these findings indicate that microbiota metabolites contribute to colonization resistance to *Cryptosporidium* infection.

---

Cryptosporidiosis, primarily caused by *Cryptosporidium hominis* or *C. parvum* in humans, manifests as self-limiting diarrhea in healthy individuals and severe, chronic diarrhea in immunocompromised patients, particularly those with HIV/AIDS^1^. Despite the parasite’s worldwide distribution^2^, the true global burden of cryptosporidiosis was underestimated until the seminal Global Enteric Multicenter Study (GEMS) unexpectedly revealed that *Cryptosporidium* is second only to rotavirus as the cause of moderate-to-severe diarrhea in infants and toddlers (0 – 24 months) in resource-poor countries^3–5^. Importantly, cryptosporidiosis in young children also increases their risk for severe malnutrition, long-term growth stunting, and death^3,6,7^. However, for reasons not well understood, the incidence of cryptosporidiosis drops dramatically in children older than two years of age in these same populations^3,4^. This inverse correlation between age and susceptibility to cryptosporidiosis is not specific to humans: other zoonotic hosts including dairy cows^8^, broiler chickens^9,10^ and mice^11,12^ are highly susceptible to *Cryptosporidium* as neonates but relatively resistant if infected as juveniles or adults. Although immune system maturation is an important factor in the development of resistance to *Cryptosporidium* infection^13,14^, dramatic changes in the composition and diversity of the gut microbiome during early development also play a pivotal role in the incidence and severity of infection^11,15,16^. For example, one prospective study of infants in Bangladesh found that the microbiota of infants that developed clinical cryptosporidiosis were less diverse one month prior to the onset of diarrheal symptoms than the microbiota of infected but asymptomatic infants^16^. Additionally, studies in adult mice have consistently shown that loss of the microbiota either through antibiotic treatment or gnotobiotic rearing sharply increases susceptibility of animals to *Cryptosporidium* infection^15,17^.

Although a diverse microbiota appears to protect against cryptosporidiosis, little is known about how the microbiota influences *Cryptosporidium* infection on a mechanistic level. One possibility is that the microbiota primes the immune system to respond to enteric pathogens. Indeed, one study in neonatal mice observed that stimulation of the intestinal immune response with poly(I:C) reduced *C. parvum* load but only when the gut microflora was present^18^. A second possibility is that metabolites in the diet or produced by the microbiota, rather than the bacteria themselves, could affect *Cryptosporidium* growth. Precedent for this idea comes from a human challenge study that found high fecal indole levels (>2.5 mM), or indole-producing bacteria, were protective against the development of diarrhea in volunteers challenged with *C. hominis* or multiple strains of *C. parvum*^19^. Additionally, prior studies have also shown that dietary supplementation with arginine can decrease severity of *C. parvum* infection in malnourished neonatal mice by stimulating the production of nitric oxide^20^.

We previously postulated that increased susceptibility to *Cryptosporidium* in neonates is partly explained by the presence of gut metabolites that enhance parasite growth^11^. Metabolites are known to undergo profound changes during development, concordant with alteration in the microbiota^11,21^. Accordingly, we focused on the potential role gut metabolites in adults may play in the development of age-dependent resistance to the parasite. We identified several classes of bacterial metabolites that inhibit parasite growth including indoles that act by impairing host mitochondrial function, leading to a reduction in total ATP levels in the host cell and reduced parasite growth in vitro and in vivo. Thus, high indole levels in the gut may decrease *Cryptosporidium* burden by depriving the parasite of essential host-derived nutrients.

## RESULTS

### Microbial metabolite screen identifies inhibitors of *C. parvum* growth in vitro

We screened a collection of 85 metabolites (Table S1) that are associated with the adult murine gut microbiota^22^ for their effects on *C. parvum* (*Cp*) infection in a human ileocecal adenocarcinoma (HCT-8) cell line (Fig. 1a). *Cp* oocysts were added simultaneously with individual metabolites to HCT-8 cell monolayers, and the numbers of parasites and host cells in each well were quantified 24 hours post infection (hpi) using an automated image-based assay^11^. In total, we identified 8 metabolites that significantly inhibited *Cp* growth and none that enhanced infection (Fig. 1a and Supplementary Table 1). These inhibitory metabolites included indole and its derivative 4-hydroxyindole (4HI), three secondary bile acids or salts (deoxycholate, deoxycholic acid, and lithocholic acid), and pyridoxal hydrochloride, a precursor of the metabolically active form of vitamin B_6_, pyridoxal-5-phosphate. Dose-response curves indicated that secondary bile acids were the most potent against *Cp* (EC_50_ < 100 μM) but had low selectivity (2.5 – 3-fold) for the parasite due to host toxicity (Fig. 1b). Pyridoxal hydrochloride was less potent against *Cp* (EC_50_ = 213.2 μM) than the secondary bile acids but had the highest fold selectivity (44.6-fold) for the parasite (Fig. 1b). Indole and 4HI were the least potent (EC_50_ = 410.9 μM and 1255 μM, respectively) but exhibited less host cell toxicity than secondary bile acids (Fig. 1b). Indole is a well-known microbial metabolite that exhibits antimicrobial activity (e.g., *Escherichia coli, Pseudomonas aeruginosa, Clostridium difficile*)^23–26^. Moreover, fecal indole levels correlate with resistance to *C. hominis* or *C. parvum* in human challenge studies^19^. Thus, we focused on defining the potential mechanism of action of indoles against *Cp*.

**Fig. 1:**
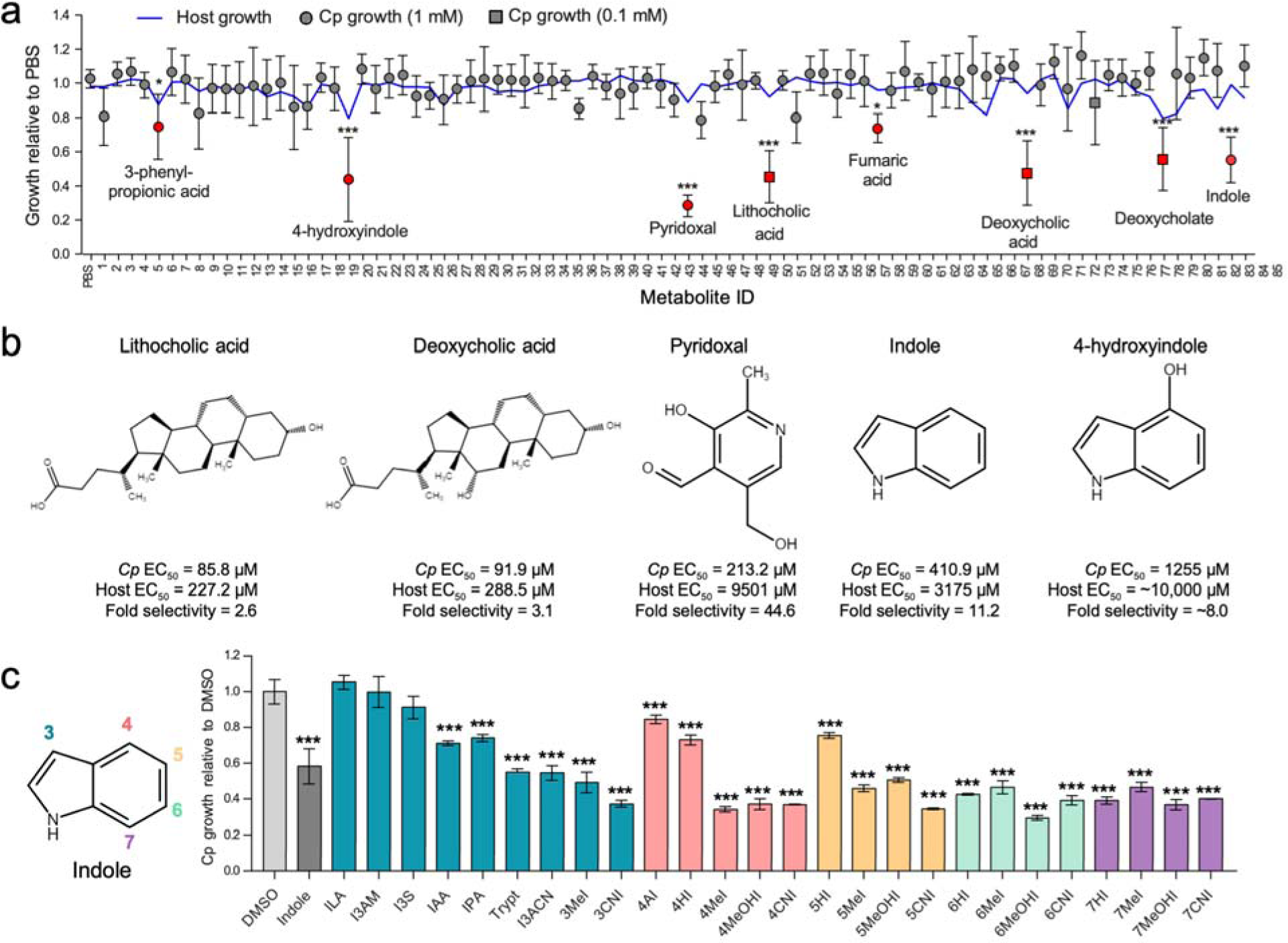
Gut metabolites, specifically secondary bile acids and indoles, inhibit *C. parvum* infection in vitro. a) Effects of 85 intestinal metabolites at 1 mM (circles) or 0.1 mM (squares) on *C. parvum* (*Cp*) infection in HCT-8 cells 24 hpi. Data plotted represents mean ± S.D. of *Cp* or mean host cell numbers (blue line) relative to PBS control for six independent experiments. Differences between *Cp* numbers for each metabolite and the PBS control were analyzed using a one-way ANOVA followed by a Dunnett’s test for multiple comparisons. Metabolites that significantly inhibited *Cp* growth are indicated in red. \**P* < 0.05, \*\*\**P* < 0.001. b) Chemical structures of five inhibitory metabolites with their respective EC_50_ values for *Cp* and host cells and fold selectivity (host EC_50_ divided by *Cp* EC­_50_). EC_50_ values were calculated using a nonlinear regression curve fit with six replicates (three technical replicates from two independent experiments) per concentration. c) Screen of indole analogs (1 mM) modified at the 3-(teal), 4-(pink), 5-(orange), 6-(green), or 7-carbon (purple) positions and their effects on *Cp* infection in HCT-8 cells. Data plotted represents mean ± S.D. of six replicates (three technical replicates from two independent experiments). Differences between mean *Cp* numbers for each metabolite and the DMSO control were analyzed using a one-way ANOVA followed by a Dunnett’s test for multiple comparisons. \*\*\**P* < 0.001. ILA = indole-3-lactic acid; I3AM = indole-3-acetamide; I3S = indoxyl-3-sulfate; IAA = indole-3-acetic acid; IPA = indole-3-propionic acid; Trypt = tryptamine; I3ACN = indole-3-acetonitrile; MeI = methylindole; CNI = cyanoindole; AI = aminoindole; HI = hydroxyindole; MeOHI = methoxyindole.

### Indole analogs modified at different carbon positions significantly inhibit *C. parvum* infection

Although both indole and 4HI significantly inhibited *Cp* growth, indole was 3X more potent than 4HI, while another indole analog included in the primary screen, 5-hydroxyindole-3-acetic acid, had no effect on *Cp* (Supplementary Table 1). These results suggest that modifications at different carbon positions along the pyrrole or benzene rings of indole (Fig. 1c) may change the potency of indole analogs. To examine the effect of modification position on indole efficacy against *Cp*, we tested a panel of 25 additional indole analogs modified at the 3-, 4-, 5-, 6- and 7-carbon positions of indole using the in vitro growth assay (Figure 1c). Interestingly, nearly all the indole analogs (22 out of 25) significantly inhibited *Cp* infection (Figure 1c and Supplementary Table 2). The three exceptions were all modified at the 3-carbon position: indole-3-lactic acid (ILA), indole-3-acetamide (I3AM), and indoxyl-3-sulfate (I3S). However, other indoles modified at the 3-carbon position, such as indole-3-propionic acid (IPA) and 3-methylindole (also called skatole), did inhibit *Cp,* indicating that the side chain composition is more important than its position on the indole molecule. Indeed, certain side chains were more potent than others, particularly methyl, methoxy and cyano groups. Of the 22 indoles that inhibited *Cp*, 11 were more potent than indole, and we chose 7-cyanoindole (7CNI) as the best analog for further study based on its combined potency and selectivity.

### Indole inhibition of *C. parvum* does not depend on the aryl hydrocarbon receptor pathway

Many metabolites produced from microbial or host tryptophan metabolism including indole and its derivatives serve as endogenous ligands for the aryl hydrocarbon receptor (AhR) in animals^27^. Furthermore, synthetic indoles such as 4-methylindole, 6-methylindole, and 7-methoxyindole have been shown to activate AhR signaling in vitro^28^. To determine whether activation of the AhR pathway by indole analogs is sufficient to inhibit parasite growth, we treated *Cp* infected cultures with agonists of AhR including tryptophan metabolites kynurenic acid^29^ and 6-formylindolo(3,2-b)carbazole FICZ^30,31^ as well as a highly specific synthetic AhR agonist, VAF347^32^. Although indole and 4MeI significantly inhibited *Cp* in a dose-dependent manner (Fig. 2a), none of the non-indole AhR agonists affected *Cp* growth (Fig. 2a), even though the expression of AhR target genes AhRR and CYP1A1 were significantly upregulated in HCT-8 cells after 24 h treatment with AhR agonists (Fig. 2b). To confirm that the AhR pathway is not involved in indole inhibition of *Cp*, we knocked out the AhR gene in HCT-8 cells using CRISPR/Cas9 to disrupt the first exon of the gene. We isolated two clonal lines, one with a single bp insertion (AhR KO 1) and the other with an 11 bp deletion (AhR KO 2) (Supplementary Fig. 1). Both mutations are predicted to cause a frame shift with premature stop codons that result in truncated proteins of 29 and 25 amino acids, respectively, (the full-length protein is 848 AA, Supplementary Fig. 1). Importantly, we confirmed that both AhR KO lines lost the ability to upregulate CYP1A1 gene expression upon VAF347 treatment (Fig. 2c). However, *Cp* remained sensitive to indole and 4HI inhibition in AhR KO cells (Fig. 2d), indicating that indoles do not act through the host AhR pathway to inhibit *Cp*.

**Fig 2.**
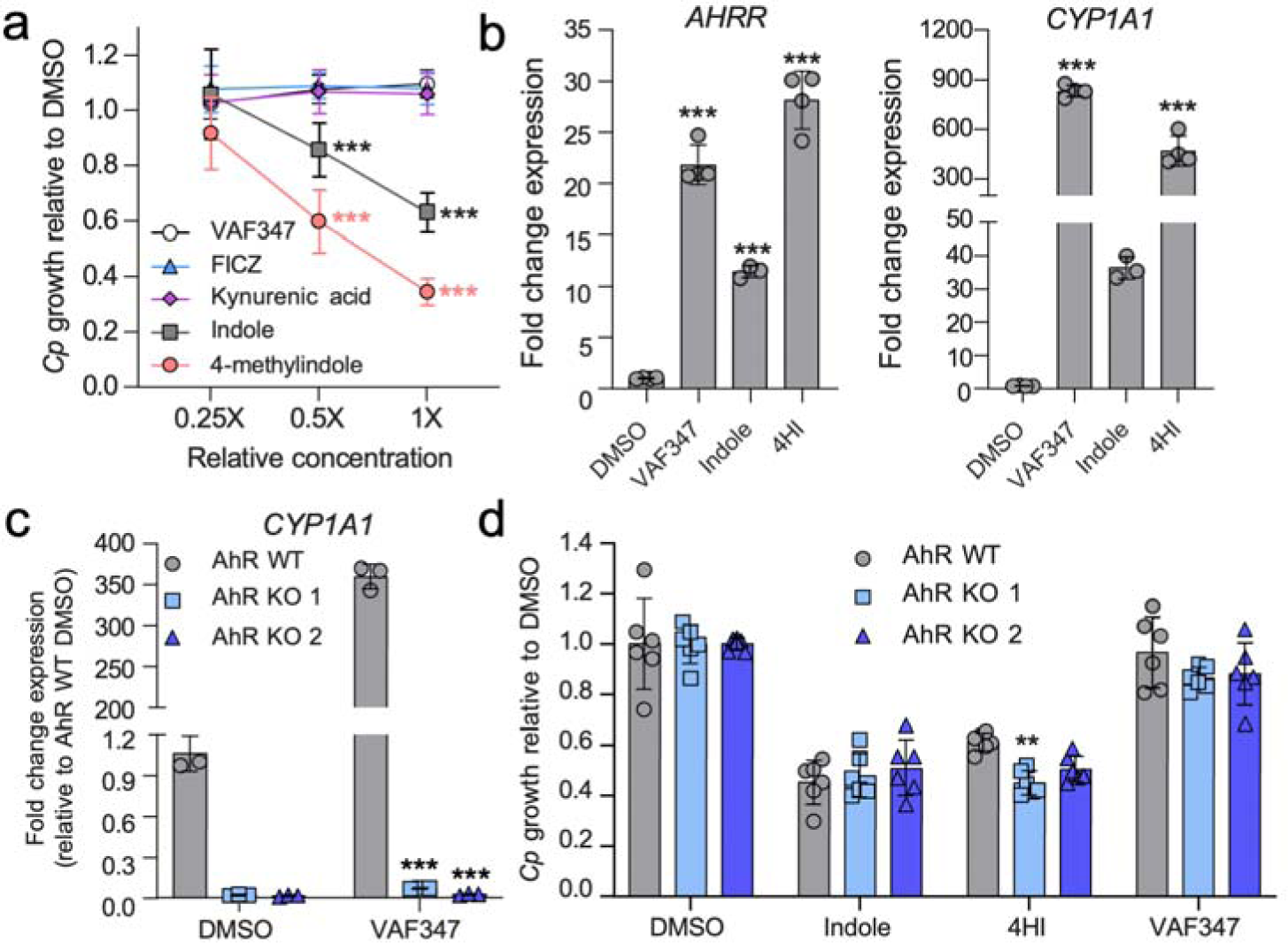
Indoles do not inhibit *C. parvum* through the host AhR pathway. a) Ratio of *C. parvum* (*Cp*) relative to DMSO control in HCT-8 cells after 24 h treatment with serial dilutions of AhR agonists. Starting concentrations (1X) were 1 μM for VAF347 and FICZ and 1 mM for kynurenic acid, indole and 4-methylindole. Data plotted represents mean ± S.D. of nine replicates (three technical replicates from three independent experiments). Differences between mean *Cp* ratio in 1X or 0.5X treated cultures versus 0.25X-treated cultures for each compound were analyzed with a two-way ANOVA followed by a Dunnett’s test for multiple comparisons. \*\*\**P* < 0.001. b) Fold change in gene expression of human *AHRR* or *CYP1A1* genes (normalized to *GAPDH)* in uninfected HCT-8 cultures after 24 h treatment with VAF347 (250 nM), indole (1.5 mM) or 4-hydroxyindole (4HI, 2.5 mM) relative to 1% DMSO control. Data plotted represents mean ± S.D. of 3 – 4 technical replicates from a single experiment. For each gene, differences between fold change in gene expression for each treatment versus DMSO control were analyzed with a one-way ANOVA followed by a Dunnett’s test for multiple comparisons. \*\*\**P* < 0.001. c) Fold change in gene expression of human *CYP1A1* genes (normalized to *GAPDH*) in uninfected HCT-8 AhR WT (gray) or KO cell lines (blue) after 24 h treatment with VAF347 (500 nM) relative to 1% DMSO control. Data plotted represents mean ± S.D. of three technical replicates from a single experiment. Differences between fold change in gene expression for each AhR KO versus AhR WT cell line for each treatment were analyzed with a two-way ANOVA followed by a Dunnett’s test for multiple comparisons. \*\*\**P* < 0.001. d) Ratio of *Cp* relative to DMSO control in HCT-8 AhR WT (gray) or KO cell lines (blue) after 24 h treatment with 0.5% DMSO, indole (1 mM), 4HI (1 mM) or VAF347 (500 nM). Data plotted represents mean ± S.D. of six replicates (three technical replicates from two independent experiments). Differences between *Cp* ratio in each AhR KO versus AhR WT cell line for each treatment were analyzed with a two-way ANOVA followed by a Dunnett’s test for multiple comparisons. \*\**P* < 0.01.

### Indoles delay intracellular life cycle progression of *C. parvum*

To test whether indoles inhibit parasite invasion and/or asexual replication, we quantified *Cp* growth in cultures treated with EC_90_ concentrations of either indole or 7CNI at time points corresponding to *Cp* invasion and parasitophorous vacuole formation (0 – 4 hpi), intracellular replication (4 – 24 hpi), or both processes (0 – 24 hpi). We also pre-treated host cells for 20 h before infection to see if indoles “prime” host cells against *Cp* infection. Pre-treatment of host cells did not affect subsequent *Cp* infection, and neither indole nor 7CNI inhibited *Cp* growth when cells were treated for the first four hours of infection only (Fig. 3a). In contrast, when treatment was started 4 hpi, indole and 7CNI significantly inhibited *Cp* infection to nearly the same extent as in cultures treated for the full 24 h (Fig. 3a). Taken together, these results indicate that 1) indoles must be present during infection to inhibit *Cp,* 2) indoles do not impair *Cp* excystation, invasion or parasitophorous vacuole formation, and 3) indoles act at some point during the intracellular replicative stages of the parasite.

**Fig. 3:**
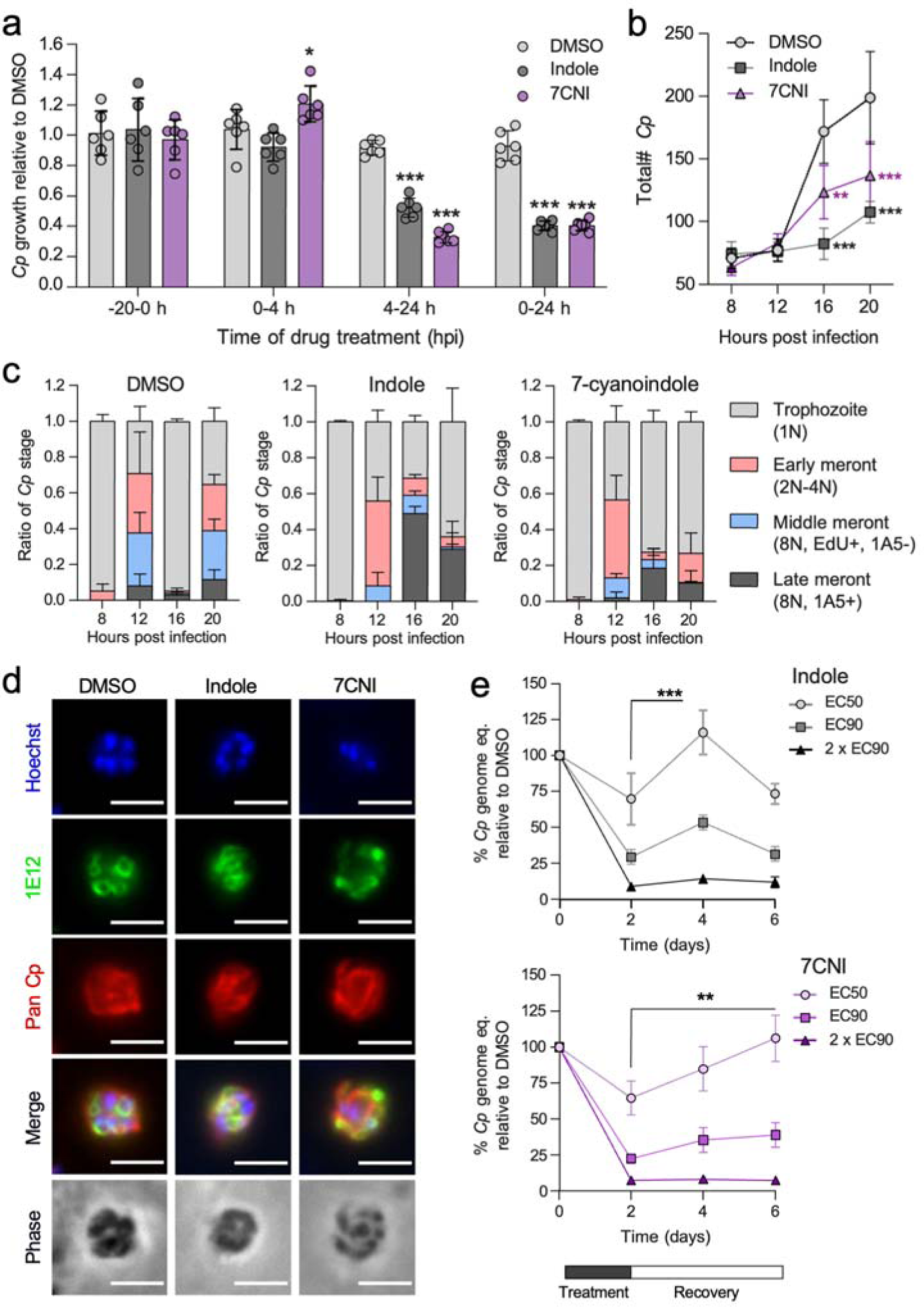
Indoles delay *C. parvum* life cycle progression. a) Ratio of *C. parvum* (*Cp*) numbers relative to DMSO control in HCT-8 cells after treatment with 1% DMSO or EC_90_ concentrations of indole (880 μM) or 7-cyanoindole (7CNI, 500 μM) for the indicated hours post infection (hpi). Data plotted represents mean ± S.D. of six replicates (three technical replicates from two independent experiments). Differences between mean *Cp* ratio in indole or 7CNI-treated cultures vs in the DMSO control at each time point were analyzed with a two-way ANOVA followed by a Dunnett’s test for multiple comparisons. \**P* < 0.05, \*\*\**P* < 0.001. b) Total number of *Cp* in HCT-8 cultures treated with 1% DMSO or EC_90_ concentrations of indole or 7CNI for the indicated hours post infection. Data plotted represents mean ± S.D. of three independent experiments (same experiments as in c). Differences between mean *Cp* numbers in indole or 7CNI-treated cultures vs in the DMSO control at each time point were analyzed with a two-way ANOVA followed by a Dunnett’s test for multiple comparisons. \*\**P* < 0.01, \*\*\**P* < 0.001. c) Ratio of the number of *Cp* in the trophozoite, early meront, middle meront or late meront stages in infected HCT-8 cultures treated with 1% DMSO or EC_90_ concentrations of indole or 7CNI at the indicated hours post infection. N = the number of nuclei per parasite. Data plotted represents mean ± S.D. of three independent experiments. d) Immunofluorescence images of *Cp* in HCT-8 cultures treated with 1% DMSO or EC_90_ concentrations of indole or 7CNI 22 hpi. Parasites are labeled with membrane marker 1E12 (green) and a general *Cp* antibody Pan Cp (red). Nuclei are stained with Hoechst. Scale bar = 3 μm. e) Washout experiments in *Cp*-infected air-liquid interface (ALI) cultures treated with 1% DMSO or indole at EC_50_ (577 μM), EC_90_ (1894 μM) or 2 x EC_90_ (3788 μM); or 7CNI at EC_50_ (379 μM), EC_90_ (688 μM) or 2 x EC_90_ (1376 μM) for 48 h before washout. *Cp* genome eq. were normalized to the DMSO control at each time point. Data plotted represents mean ± S.D. of six replicates (three technical replicates from two independent experiments). Differences between mean % *Cp* after washout versus mean % *Cp* at time of washout (2 dpi) for each indole concentration were analyzed using a two-way ANOVA followed by a Dunnett’s test for multiple comparisons. \*\**P* < 0.01, \*\*\**P* < 0.001.

After invasion, *Cp* begins its intracellular asexual cycle as a single nucleus trophozoite that undergoes three rounds of DNA replication with incomplete cytokinesis (early and middle meronts), followed by separation of the individual nuclei and membrane engulfment to form eight mature type I merozoites (late meront)^33^. The mature merozoites then egress from the host cell and invade a neighboring host cell to restart the asexual cycle. To more precisely define when indoles act during the intracellular stages of the parasite, we treated infected cultures with indole or 7CNI at EC_90­_ concentrations while adding the thymidine analog EdU in 4 h pulses to label replicating parasite DNA and also staining for stage specific markers, as described previously^33–35^. We then quantified the total number of parasites and the ratio of intracellular life cycle stages present at each time point (Supplementary Fig. 2). We found that the total number of *Cp* did not significantly differ between DMSO and indole or 7CNI-treated cultures until the start of the second round of merogony (∼16 hpi, Fig. 3b), indicating that indoles inhibit parasite development. When the progression of specific replicative stages was examined over the first 24 hpi, it was evident that indole and 7CNI delayed progression through sequential life cycle stages, rather than stalling development at a specific time point (Fig 3c). To determine if indoles prevent merozoite maturation, egress, and/or re-invasion similar to the previously studied compound KDU691^33^, we treated infected cultures with EC_90_ indole or 7CNI for 22 h before fixing and labeling with 1E12, a monoclonal antibody that labels the parasite surface membrane^35^. We did not observe any obvious morphological defects in mature merozoites in indole or 7CNI-treated cultures (Fig. 3d), indicating indoles do not prevent formation of mature merozoites.

### Indole inhibition is partly reversible after washout in a long-term culture system for *C. parvum*

To examine the reversibility of treatment, we employed an air-liquid interface (ALI) transwell culture system that supports both asexual and sexual development and allows long-term growth of *C. parvum*^34,36^. We treated infected cultures with indole or 7CNI at EC_50,_ EC_90_ and 2 x EC_90_ concentrations for two days before washing out the compounds and allowing the cultures to recover for 2 – 4 days before quantification of *Cp* and host genomes. After treatment with indole or 7CNI EC_50_ concentrations, there was significant recovery in *Cp* growth 2 days after indole washout and 4 days after 7CNI washout (Fig. 3e). Some growth recovery (though not significant) occurred after washout of cultures treated with EC_90_ indole or 7CNI, while no recovery was observed in cultures treated with 2 x EC_90_ indole or 7CNI (Fig. 3e). However, the lack of parasite recovery at the highest concentrations of indole and 7CNI was likely affected by the high level of host toxicity at these concentrations (Supplementary Fig. 3). Thus, inhibition of *Cp* growth by indoles is partially reversible at lower concentrations when host cell viability is not affected. These results suggest that, while indoles impair growth when present, there is rapid restoration of an environment that is favorable for parasite development upon removal.

### Indole treatment upregulates host genes involved in ER stress and membrane transport

To determine if indoles alter host gene expression to create an unfavorable host cell environment for *Cp* growth, we performed RNA-seq on HCT-8 cells treated with *Cp* EC_90_ concentration of indole or 1% DMSO for 4 or 12 h. When comparing all indole-treated (N = 6) with all DMSO-treated (N = 5) samples using length of treatment as a co-variable, we identified 68 genes that were significantly differentially expressed after indole treatment (FDR *P* value < 0.05, absolute fold change > 2): 57 upregulated genes (including AhR target gene *CYP1A1*) and 11 downregulated genes (Fig. 4a, Supplementary Table 3). Hierarchical clustering analysis of differentially expressed genes revealed that gene expression patterns separated into 3 main clusters: downregulated after indole treatment, highly upregulated after 4 h of indole treatment, and highly upregulated after 12 h of indole treatment (top 30 genes are shown in Fig. 4b, all genes listed in Supplementary Table 3). Gene Ontology (GO) process analysis performed in Enrichr on the subset of genes upregulated after 4 h of indole treatment found that the two most significant pathways were in response to ER stress (GO:0034976) and apoptotic signaling in response to ER stress (GO:0070059, Fig. 4c). Interestingly, the expression of genes associated with these pathways, namely *DDIT3, DNAJB9,* and *CHAC1*, goes down with longer exposure to indole (Fig. 4b). In contrast, five out of the top 10 pathways upregulated after 12 h of indole treatment are involved in membrane transport of carboxylic acids (GO:0046942), monocarboxylic acids (GO:0015718), amino acids (GO:0015804 and GO:0006865), or nitrogen compounds (GO:0071705, Fig. 4d). Taken together, these transcriptomics data suggest that short treatment with indole induces an ER stress response, which may cause the cells to upregulate transporters to restore imbalances in important nutrients. We reasoned that indole may be competing with the import of tryptophan or other aromatic amino acids like phenylalanine, both essential amino acids that cannot be made by the host cell. However, supplementation of cell culture medium with additional tryptophan or tryptophan plus phenylalanine did not rescue indole inhibition of *Cp* growth in HCT-8 cells (Fig. 4e). Thus, indole inhibition of *Cp* is likely not due to a tryptophan deficiency in the host but may still result from a lack of another essential nutrient.

**Fig 4.**
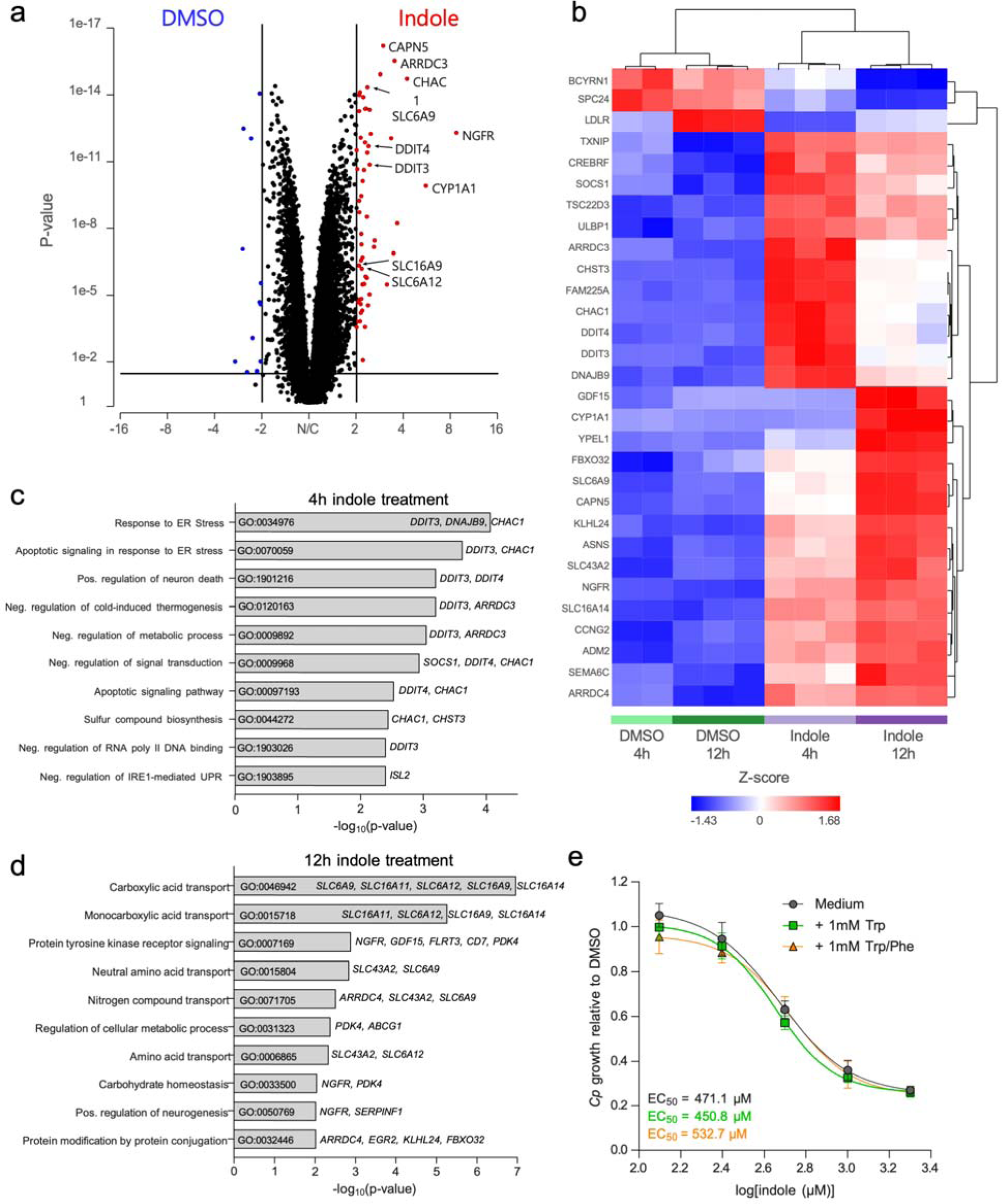
Indole induces ER stress and transporter upregulation in HCT-8 cells. a) Volcano plot of fold-change vs *P-*value after GSA analysis of indole vs DMSO-treated HCT-8 cells highlighting genes significantly (*P* < 0.05) upregulated (red) or downregulated (blue) after indole treatment by >2-fold. b) Hierarchical clustering analysis of the 30 most differentially regulated genes (FDR-corrected *P* < 1×10^−9^) between indole and DMSO-treated HCT-8 cells. c,d) Gene Ontology (GO) pathway analysis performed in Enrichr using genes significantly upregulated after c) 4 h or d) 12 h of indole treatment as input. Upregulated genes associated with each pathway are listed to the right of the bar graph. e) Ratio of *C. parvum* (*Cp*) relative to DMSO control in HCT-8 cells after 24 h treatment with a serial dilution of indole in growth medium supplemented with 1 mM tryptophan (Trp) or 1 mM Trp plus 1 mM phenylalanine (PHE). EC_50_ values were calculated for each medium using a nonlinear regression curve fit with six replicates (three technical replicates from two independent experiments) per indole concentration.

### Indoles impair host mitochondrial function and reduce ATP production

Previous studies have reported that indole can uncouple oxidative phosphorylation (OxPhos)^37,38^, inhibit the electron transport chain^37^, and/or decrease ATP levels^26,37,39^ in *Pseudomonas putida*^26^, rat liver^37,38^, and human enteroendocrine cells^39^. To test whether indoles inhibit mitochondrial function in HCT-8 cells, we measured the oxygen consumption rate (OCR) and respiratory capacity of cells treated with indole or 7CNI. We found that indole and 7CNI significantly reduced both basal levels and maximal levels of mitochondrial respiration in HCT-8 cells in a dose-dependent manner (Fig. 5a and Supplementary Fig. 4). As a result, the spare respiratory capacity (maximal minus basal respiratory rate) of indole and 7CNI-treated cells was also diminished (Fig. 5a and Supplementary Fig. 4), indicating that the mitochondria in these cells are less able to respond to a sudden increase in energy demand (simulated by the addition of proton ionophore and OxPhos uncoupler FCCP in the assay). This dose-dependent impairment of mitochondrial function by indole and 7CNI translated into a significant reduction in ATP production rate by the mitochondria (Fig. 5b and Supplementary Fig. 4). Although ATP production rates by glycolysis remained similar between all treatments, there was a lower total ATP production rate in indole or 7CNI-treated cells (Fig. 5b and Supplementary Fig. 4).

**Fig 5.**
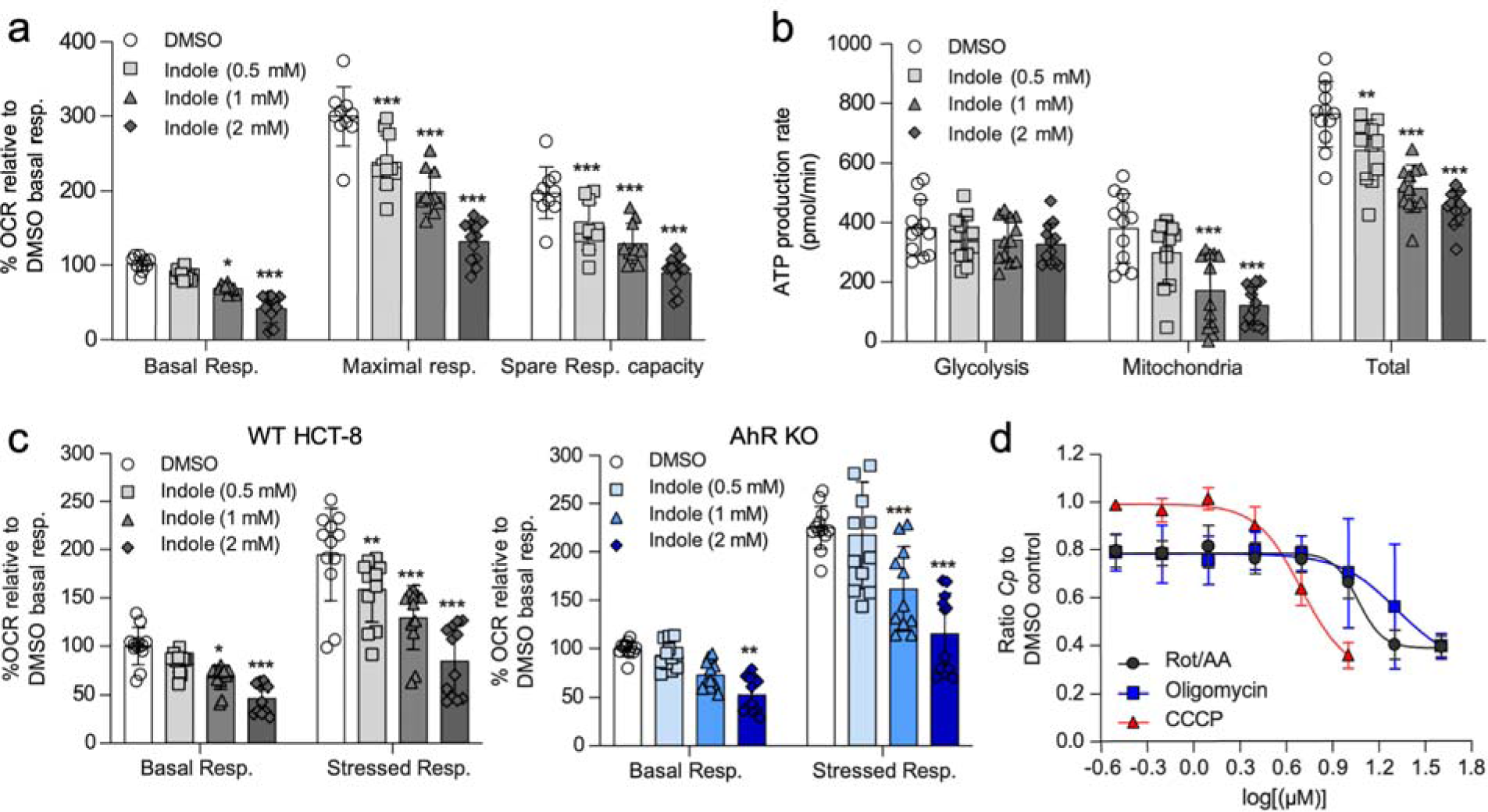
Indole impairs host mitochondrial ATP production. a) Metabolic analysis using the Seahorse XF Cell Mito Stress Test kit on HCT-8 cells treated for 18 h with 1% DMSO or indole (0.5 mM, 1 mM or 2 mM). Data calculated as a percentage of the oxygen consumption rate (OCR) for each well relative to the mean basal OCR of DMSO control cells for that experiment. Spare respiratory capacity = maximal respiratory rate – basal respiratory rate for each well. Data plotted represents mean ± S.D. of 12 replicates (six technical replicates from two independent experiments). Differences between *%* OCR for each indole concentration vs the DMSO control for each measurement were analyzed with a two-way ANOVA followed by a Dunnett’s test for multiple comparisons. \**P* < 0.05, \*\*\**P* < 0.001. b) Metabolic analysis using the Seahorse XF Real-time ATP Rate assay on HCT-8 cells treated for 18 h with 1% DMSO or indole (0.5 mM, 1 mM or 2 mM). Data plotted represents mean ± S.D. of ATP production rate (pmol/min) produced by glycolysis, the mitochondria, or total ATP (glycolysis + mitochondrial ATP rates) for 12 replicates (six technical replicates from two independent experiments). For each source of ATP, differences between ATP production rate for each indole concentration vs the DMSO control were analyzed with a two-way ANOVA followed by a Dunnett’s test for multiple comparisons. \*\**P* < 0.01, \*\*\**P* < 0.001. c) Metabolic analysis using the Seahorse XF Cell Energy Phenotype Test kit on HCT-8 AhR WT cells (gray) or AhR KO cells (blue) treated for 18 h with 1% DMSO or indole (0.5 mM, 1 mM or 2 mM). Data calculated as a percentage of OCR for each well relative to the mean basal OCR of DMSO control cells for that experiment. Data plotted represents mean ± S.D. of 12 replicates (six technical replicates from two independent experiments). For each cell line, differences between *%* OCR for each indole concentration vs the DMSO control were analyzed with a one-way ANOVA followed by a Dunnett’s test for multiple comparisons. \**P* < 0.05, \*\**P* < 0.01, \*\*\**P* < 0.001. d) Ratio of *C. parvum* (*Cp*) relative to DMSO control in HCT-8 cells after 24 h treatment with serial dilutions of mitochondrial Complex I and III inhibitors rotenone and antimycin A, respectively (Rot/AA); ATP synthase inhibitor oligomycin; or proton gradient uncoupler carbonyl cyanide m-chlorophenyl hydrazone (CCCP). Inhibition curves were calculated for each compound using a nonlinear regression curve fit with six replicates (three technical replicates from two independent experiments) per concentration.

Unlike its host cell, *Cp* does not contain an intact mitochondrion but instead has a remnant mitosome organelle that lacks both a functional TCA cycle and a cytochrome-dependent electron transport pathway and therefore does not directly contribute to ATP production^40^. Despite lacking a conventional electron transport chain, *Cp* mitosomes do sequester cationic dyes traditionally used to stain live mitochondria, such as Mitotracker CMXRos, suggesting the mitosome has an alternative mechanism to generate a membrane potential^41^. Here, we verified that a potent depolarizer of mitochondrial membrane potential, CCCP, also decreases Mitotracker staining in the *Cp* mitosome (Supplementary Fig. 5). Similarly, treatment of *Cp* with indole or 7CNI, whether briefly (e.g., 2 h) or for an extended time period (e.g., 24 h), significantly decreased Mitotracker staining of *Cp* mitosomes, indicating indoles likely impair the membrane potential of this organelle (Supplementary Fig. 5).

Although indole inhibition of *Cp* does not appear to act through the AhR pathway (Fig. 2), AhR responsive genes were clearly upregulated in the RNA-seq experiments (Fig. 4) raising the possibility that this pathway might affect mitochondrial function. To test this possibility, we examined mitochondrial function using the HCT-8 AhR KO 1 cell line. We found that mitochondrial basal respiratory rate and stressed respiratory rate (measured after the simultaneous addition of FCCP and oligomycin, an ATP synthase inhibitor) were significantly impaired in a dose-dependent manner in both HCT-8 WT and AhR KO cells (Fig. 5c). Thus, impaired mitochondrial respiration is associated with inhibition of *Cp* growth in a pathway that does not depend on AhR. To determine whether inhibiting mitochondrial function was sufficient to inhibit *Cp* growth independently of indole, we performed the *Cp* growth assay in HCT-8 cells after treating with serial dilutions of OxPhos inhibitors oligomycin, CCCP (proton ionophore structurally similar to FCCP), or a combination of rotenone and antimycin A (Rot/AA), inhibitors of Complex I and Complex III, respectively. All three treatments inhibited *Cp* growth in a dose-dependent manner with EC_50_ values of 11.5 μM for Rot/AA, 20.6 μM for oligomycin, and 5.0 μM for CCCP (Fig. 5d), levels that were not due to host cell toxicity (Supplementary Fig 4).

### Oral 7-treatment with cyanoindole treatment or introduction of indole-producing microbes temporarily suppresses *C. parvum* infection *in vivo*

To determine whether oral treatment with indoles could alter infection in a mouse model of cryptosporidiosis, we treated *Cp-*infected interferon gamma knock out (GKO) mice with 50 mg/kg indole or 7CNI by oral gavage twice daily for 7 days starting on the day of infection (Fig. 6a). GKO mice are naturally susceptible to infection and when infected with the AUCP-1 strain of *Cp* used here, they do not require antibiotic treatment to result in a robust infection characterized by shedding of high numbers of oocysts. Although indole had no effect on *Cp* infection *in vivo*, 7CNI treatment significantly decreased the number of *Cp* oocysts shed in the feces of mice by 5 dpi (Fig. 6b). The mean number of *Cp* oocysts per mg feces remained lower in 7CNI-treated mice at 7 dpi (the last day of treatment) but rebounded by 9 dpi (Fig. 6b). Similarly, 7CNI-treated mice weighed significantly more than vehicle- or indole-treated mice 5 and 7 dpi but started to lose weight rapidly after treatment was terminated (Fig 6c). Although 7CNI treatment did not significantly increase survival rate, it did prolong the median survival time to 23.5 dpi compared to 11 dpi or 17 dpi for vehicle or indole-treated mice, respectively (Fig 6d).

**Fig 6.**
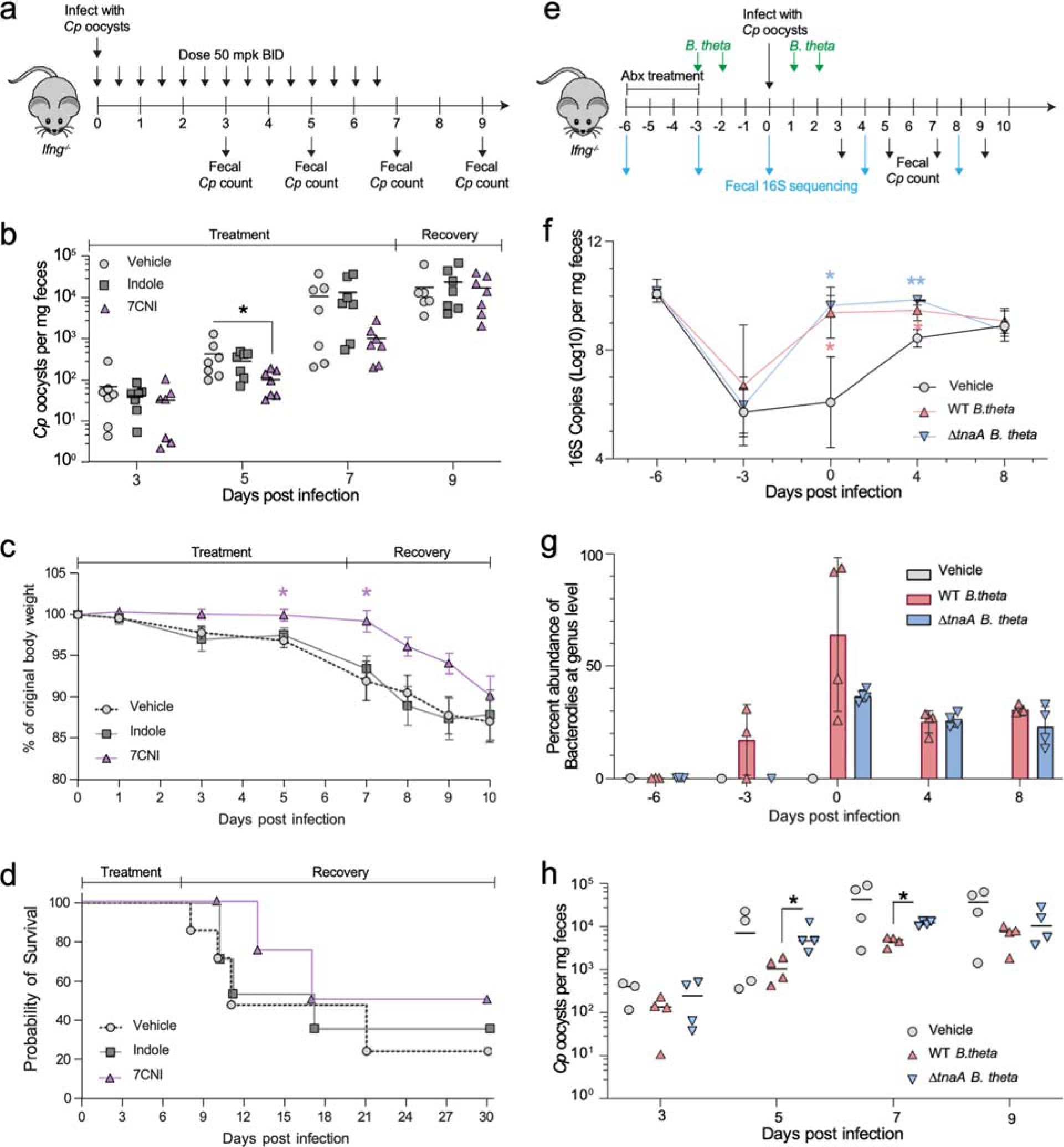
Exogenous indole treatment, or reconstitution with indole-producing bacteria, suppresses *C. parvum* infection in GKO mice. a) GKO mice were treated twice daily by gavage with vehicle (10% DMSO in water), or 50 mg/kg indole or 7-cyanoindole (7CNI) for 7 days. *Cp* oocysts numbers were quantified from a single fecal pellet per mouse collected 3, 5, 7 and 9 dpi. All data plotted represents 7 mice per treatment group sampled over time from two independent experiments. b) *Cp* oocysts per mg feces for each mouse at the indicated dpi. Statistical analyses comparing each treatment group to vehicle control on individual days were performed using two-tailed Mann-Whitney U tests. \**P* < 0.05. c) Percent of original body weight plotted as mean ± S.D. Statistical analysis performed using a mixed-effects model with a Geisser-Greenhouse correction for matched values, followed by a Dunnett’s test for multiple comparisons. \**P* < 0.05. d) Combined survival curves of all 7 mice for the first 10 days then for the 4 mice from the second experiment for days 11 – 30. e) GKO mice were treated with antibiotic to suppress endogenous flora and then reconstituted with WT *B. theta* or *ΔtnaA B. theta* followed by challenge with *Cp*. Oocysts numbers were quantified from fecal pellets collected 3, 5, 7 and 9 dpi. All data plotted represents 4 mice per treatment group sampled over time. f) Estimation of bacterial burdens in the gut by 16S rRNA qPCR. Means ± S.D. Statistical analyses comparing vehicle to WT *B. theta* group (shown in blue asterisk) or *ΔtnaA B. theta* group (shown in red asterisk) on individual days were analyzed with a two-way ANOVA followed by a Dunnett’s test for multiple comparisons. \**P* < 0.05, \*\**P* < 0.01. g) Relative abundance of Bacteroides at the genus level from each mouse at the indicated dpi. h) *Cp* oocysts per mg feces for each mouse at the indicated dpi. Statistical analyses comparing WT *B. theta* group to *ΔtnaA B. theta* group on individual days were performed using two-tailed Mann-Whitney U tests. \**P* < 0.05.

To extend the studies on the role of indole in protection in vivo, we eliminated the endogenous microflora by treatment with antibiotics and then reconstituted the microbiota with indole-producing bacteria. A variety of gram-negative bacteria express the tryptophanase gene (TnaA), which converts tryptophan to indole ^42^. Among these, members of the Bacteroidales and Clostridiales orders are among the most abundant taxa with the ability to generate indole within the gut ^42^. Hence, we chose *Bacteroides thetaiotaomicron* (*B. theta*) for reconstitution as previous studies have shown that indoles produced by this species alter susceptibility to enterohemorrhagic *E. coli* infection by modulating virulence gene expression ^43^. For these experiments, mice were treated with a cocktail of antibiotics (Abx) to suppress the endogenous flora and then reconstituted by oral gavage with wild type *B. theta* that expresses TnaA or a mutant that lacks this biosynthetic pathway (ΔtnaA) (Fig 6e). Following antibiotic treatment, the microbiota was depleted as assessed by qPCR analyses of 16S copy numbers (Fig. 6f). Reconstitution with *B. theta* led to a rapid recovery of total bacterial levels, while vehicle-treated control mice returned more gradually to pre-antibiotic levels of gut bacteria (Fig. 6f). As expected, the richness of the endogenous bacterial communities was decreased by Abx treatment, and the number of observed species remained low in mice colonized with *B. theta* or administered vehicle (Supplemental Fig. 6). For all mice, the bacterial community structure, analyzed by weighted UniFrac distance, was disrupted by Abx, with the communities of mice administered WT or mutant *B. theta* converging by 0 dpi while those of vehicle-treated mice remained disrupted and returned to a pre-antibiotic structure by 8 dpi (Supplemental Fig. 6). When analyzed taxonomically, reconstitution led to robust levels of Bacteroidales by 0 dpi in mice given wild type (WT) or mutant *B. theta*, which are absent in vehicle-treated mice (Supplemental Fig. 7). Analysis of these data at the genus level indicated that although *Bacteroides* was generally undetected prior to Abx, it became a dominant group in mice administered WT or mutant *B. theta,* with the only amplicon sequence variant identified for *Bacteroides* originating from *B. theta* (Fig. 6g). In contrast, in mice given the vehicle control, *Bacteroidales* and *Bacteroides* were absent (Supplementary Fig. 7, Fig. 6g). Having established that Abx treatment and reconstitution was effective in replacing the endogenous flora with WT *B. theta* capable of producing indole and the tnaA mutant that lacks this ability, we tested the reconstituted nice for susceptibility to infection. Reconstitution with WT *B. theta* led to a significant reduction in oocyst shedding at 5 and 7 dpi, when compared to mice reconstituted with the ΔtnaA mutant, supporting a role for indole production in resistance to infection (Fig. 6h). The difference in shedding was transient and the eventual convergence in susceptibility in mice given WT and mutant bacteria may be due to the emergence of other taxa, notably Clostridiales (Supplemental Fig. 7), members of which are also known to produce indoles ^42^.

## DISCUSSION

As an enteric pathogen, *Cryptosporidium* parasites are exposed to the billions of microbial inhabitants of the gut as well as the vast array of molecules they produce. Our screen of adult murine gut metabolites identified indole and its derivative, 4-hydroxyindole, as inhibitors of *C. parvum* growth in human cells. Further testing extended these findings to show that most indole analogs can inhibit the parasite regardless of modification position or side chain composition. Although indoles are known to signal through AhR, KO cell lines lacking AhR revealed that indole restriction of *C. parvum* is independent of this pathway. Instead, we found that indoles likely impact *C. parvum* growth indirectly by suppressing host mitochondrial respiration, leading to decreased ATP levels. Indole treatment also resulted in loss of membrane potential in the parasite mitosome, suggesting indoles can also impair function of this organelle. Indole treatment delayed progression of the parasite life cycle and was partially reversible upon washout, suggesting the effect is largely static. Treatment with indoles *in vivo* partially protected mice from *C. parvum* infection, as did reconstitution with indole-producing bacteria, suggesting that supplementation of indoles, or enhancement of microbial communities that produce them in large amounts, may offer a novel means of reducing infection.

*Cryptosporidium* spp. have severely reduced metabolic capabilities and most lack both conventional mitochondria and the enzymes necessary for the tricarboxylic acid (TCA) cycle and oxidative phosphorylation^44^. As such, *C. parvum* relies almost entirely upon host cells for their energy needs and prior studies have suggested that they may import phosphorylated nucleotides^45^, including ATP. Furthermore, several metabolomic and proteomic studies have found that *C. parvum* and *C. hominis* infections induce host mitochondrial and glycolytic activity in cell lines (HCT-8 and COLO-680N) and experimental mouse models (C57BL/6J and BALB/c)^46–49^, with a concordant increase in cellular ATP levels^46,49^. Our findings demonstrate that indoles inhibit host mitochondrial respiration in HCT-8 cells leading to a reduction in ATP production. Previous studies using intestinal enteroendocrine cells found that that indoles increased cellular NADH/NAD^+^ ratios, which would decrease the proton gradient across the inner mitochondrial membrane, reduce the oxygen consumption rate (OCR), and inhibit ATP synthesis^39 50^. Consistent with this model, we found that treatment with known mitochondrial ETC inhibitors rotenone, antimycin A, oligomycin and CCCP all had deleterious, dose-dependent effects on *C. parvum* growth. This mechanism of action is consistent with the observed developmental delay in *C. parvum* growth when cells were treated with indoles and with the observation that the effect was partially reversible upon washout of the compounds. In contrast, we found no evidence for a direct effect of indoles on the parasite mitosome or its uptake of related aromatic amino acids such as tryptophan. Although our findings do not rule out a potential effect directly on the parasite, available data support an indirect role for indole through inhibition of host mitochondrial respiration and reduced ATP.

Our findings also indicate that indoles directly affect the membrane potential of the mitosome, a degenerate organelle that lacks many features of normal mitochondria including an absence of DNA, complexes 2,3, and 4 of the electron transport chain, and nearly all of the enzymes of the TCA cycle^40,51^. As such, the mitosome does not generate ATP, although it maintains a membrane potential and consequently accumulates dyes like Mitotracker CMXRos. Previous studies have demonstrated that Fe-S cluster synthesis proteins are transported to the mitosome^52^, and it is also likely the site of ubiquinone biosynthesis^40,51^. The mitosome also contains an alternative oxidase that may function with NAD(P) transhydrogenase and NDH2 that likely act as an alternative proton pump to establish a membrane gradient^40,51^. Our studies demonstrate that treatment with indoles, or CCCP, results in decreased staining by Mitotracker, suggesting these agents disrupt the membrane potential and impair mitosome function, potentially contributing to the developmental delay of the parasite.

Our studies reveal that several classes of microbial metabolites impair growth of *C. parvum in vitro* including bile salts, indoles, and several short chain fatty acids (i.e. fumaric, propionic). Notably, the active metabolites identified here are characteristic of the adult microbiota and are largely absent in neonatal mice, which are highly susceptible to cryptosporidiosis^11^. Hence, the changes that occur in microbial metabolites with maturation of the microbiome likely contribute to colonization resistance seen in adults. We tested the tole of indole derivatives in vivo and demonstrated that either direct administration or reconstitution with indole-producing bacteria provide partial protection against infection. The action of indoles in vivo could arise from the observed growth inhibitor properties we measured in vitro (AhR-independent) but may also affect AhR signaling that increases barrier function and immunity in vivo ^27,53,54^. Consistent with a protective role for indole, previous studies have shown that healthy adults with high levels of fecal indole are resistant to challenge with *Cryptosporidium*^19^. Indoles are produced from dietary tryptophan by a wide range of bacterial taxa that express the TnaA gene encoding tryptophanase including Clostridia, Bacteroidia, and Gammaproteobacteria ^42^. As such, exogenously delivered metabolites such as indoles, or enhancement of bacterial communities producing such entities, might provide an adjunct to antibiotic therapy in control of cryptosporidiosis.

## MATERIALS AND METHODS

### Parasite culture and infection

*C. parvum* isolate AUCP-1 was maintained by repeated passage in male Holstein calves and purified from fecal material by sieve filtration, Sheather’s sugar flotation, and discontinuous sucrose gradient centrifugation as previously described ^55,56^. All calf procedures were approved by the Institutional Animal Care and Use Committee (IACUC) at the University of Illinois at Urbana-Champaign. Purified oocysts were stored at 4 °C in phosphate-buffered saline (PBS) plus 50 mM Tris and 10 mM EDTA (pH 7.2) for up to six months after fecal collection. Before infection of cell culture or animal models, *C. parvum* oocysts were treated in a 40% bleach solution (commercial bleach containing 8.25% sodium hypochlorite) diluted in Dulbecco’s phosphate-buffered saline (DPBS; Corning) for 10 mins on ice, then washed three times in DBPS containing 1% (wt/vol) bovine serum albumin (BSA; Sigma). Bleached oocysts were stored for up to 1 week in DBPS plus 1% BSA at 4 °C before infection. For most experiments, cells were infected with bleached oocysts added directly to the cell culture medium. For experiments requiring infection with sporozoites, bleached oocysts were excysted in 0.75% sodium taurocholate diluted in DPBS for 37 °C for 1 h, then centrifuged at 1,250 x g for 3 min to pellet the sporozoites and remove the sodium taurocholate before resuspending in appropriate cell culture medium.

### Cell lines and culture

All cell lines were cultured at 37 °C in a 5% CO_2_ incubator under normal atmospheric oxygen conditions. Human ileocecal adenocarcinoma cells (HCT-8; ATCC CCL-244) and HCT-8 AhR knock-out cell lines (this study) were maintained in RPMI 1640 ATCC Modification medium (Thermo Fisher A14091-01) supplemented with 10% fetal bovine serum (Gibco). NIH/3T3 embryonic mouse fibroblast cells (ATCC CRL-1658) were maintained in Dulbecco’s Modified Eagle’s Medium (DMEM high glucose; Sigma D6329) supplemented with 10% fetal bovine serum (Sigma). For transwell experiments, primary ileal intestinal epithelial stem cells (IECs) isolated from female C57BL/6 mice were expanded and maintained as 3D spheroid cultures in Matrigel (Corning) and 50% L-WRN conditioned medium (CM) containing 10 μM Y-27632 ROCK inhibitor (Torcis Biosciences), as previously described ^57,58^. All cell lines were confirmed to be mycoplasma-free with the e-Myco plus *Mycoplasma* PCR detection kit (Boca Scientific).

### Bacteria lines and culture

The wild type (WT) and tryptophanase knockout (*ΔtnaA*) *Bacteroides thetaiotaomicron* (*B. theta*) VPI-5482 strains were gifts from Vanessa Sperandio (University of Texas Southwestern Medical Center, Dallas, Texas, USA). Following growth on BHI-blood agar plates, single colonies were grown overnight at 37 °C in anaerobic conditions in chopped meat media. The bacteria strains were aliquoted and frozen at −80 °C in 20% glycerol until further processing. Colony forming units (CFUs) of frozen stocks were calculated based on serial dilution spotting on TSA + 5% sheep’s blood plates (Thermo Fisher Scientific).

### Animal studies

Ifng knockout mice (referred to as GKO) were purchased from Jackson Laboratories (B6.129S7-Ifng^tm1Ts/J^) or bred in house in a specific-pathogen-free animal facility on a 12:12 light-dark cycle. Mice received irradiated laboratory rodent chow (Purina 5053) and autoclaved water *ad libitum*. Mice were co-housed with siblings of the same sex throughout the experiments. Animals that became non-ambulatory during the course of infection were humanely euthanized in a SMARTBOX Auto CO_2_ euthanasia chamber. All mouse studies were approved by the Animal Studies Committee at the School of Medicine, Washington University in St. Louis.

### Initial screen of gut metabolites on *C. parvum* growth

Metabolites (Sigma-Aldrich) were dissolved in appropriate solvent (Table S1) and screened at a final concentration of 1 mM. Metabolites that displayed significant host toxicity at 1 mM were re-screened at lower concentrations, specifically 0.1 mM for deoxycholic acid (DCA), sodium deoxycholate (SDC), and lithocholic acid (LCA) and 0.05 mM for prostaglandin E2. For the inhibition assay, HCT-8 cells were plated at 1.5 x 10^5^ cells per well in black-sided, optically clear-bottomed 96-well plates (Greiner Bio-One) and grown for ∼24h until confluent. Cells were then infected with 5 x 10^5^ bleached *C. parvum* oocysts per well along with a single metabolite diluted in culture medium. After 24h of infection/treatment, cells were fixed in a 9:1 methanol:acetate solution for 5 min, washed twice with DPBS, permeabilized in 0.05% saponin in PBS for 10 min, and blocked in 0.05% saponin, 5% normal goat serum (NGS), and 5% fetal bovine serum (FBS) in PBS for 10 mins before antibody staining. Parasites were labeled with an anti-RH antibody (rabbit polyclonal antibody raised against *Toxoplasma* strain RH that also recognizes all intracellular *C. parvum* stages^35^) diluted 1:1000 in PBS containing 1% NGS and 0.1% saponin, followed by Alexa Fluor goat anti-rabbit 594 secondary antibody (Thermo Fisher Scientific, diluted same as primary antibody). Host cells were stained with Hoechst 33342 (1 μg/ml, Thermo Fisher Scientific) for 10 min. Plates were imaged with a 10X objective on a BioTek Cytation 3 cell imager (9 images per well in a 3 x 3 grid). Gen5 software version 3.02 was used to quantify the total number of parasites (puncta in the Texas Red channel) and host cells (nuclei in the DAPI channel) per well. Each metabolite was screened in triplicate in six separate experiments. Relative parasite and host cell growth were calculated as a ratio of the number of *C. parvum* or host cells, respectively, in treated vs PBS negative control wells on each plate.

### *C. parvum* growth assay

All *C. parvum* growth assays (aside from the initial metabolite screen) were performed as detailed here, with any modifications stated in the specific experimental sections. HCT-8 cells were plated at 1.5 x 10^5^ cells per well in black-sided, optically clear-bottomed 96-well plates (Greiner Bio-One) and grown for ∼24h until confluent. Cells were infected with 1 x 10^5^ bleached oocysts per well. After ∼24h of infection/treatment, cells were fixed in 4% formaldehyde for 10 min, washed twice in DPBS, and then permeabilized and blocked in PBS containing 0.1% Triton-X and 1% bovine serum albumin (BSA) for 20 mins before antibody staining. *C. parvum* parasites were labeled with Pan-Cp (rabbit polyclonal antibody raised against *C. parvum* that recognizes all stages of the parasite^34^) diluted 1:2,000 in PBS containing 0.1% Triton-X and 1% BSA, followed by Alexa Fluor goat anti-rabbit 488 secondary antibody (Thermo Fisher Scientific, diluted same as primary antibody). Host cell nuclei were stained with Hoechst 33342 (5 μg/ml, Thermo Fisher Scientific) for 20 min. Plates were imaged with a 10X objective on a BioTek Cytation 3 cell imager (9 images per well in a 3 x 3 grid). Gen5 software version 3.08 was used to quantify the total number of parasites (puncta in the GFP channel) and host cells (nuclei in the DAPI channel) per well.

### Indole analog screen

Indole analogs (Table S2) were ordered from Sigma-Aldrich (St. Louis, MO) or AA Blocks, Inc. (San Diego, CA) and reconstituted at 100 mM or 200 mM in DMSO. *C. parvum* growth assays were performed as described above with all analogs diluted to 1 mM in culture medium. Relative parasite number was calculated as a ratio of the number of *C. parvum* in treated wells vs the mean number of parasites in 1% DMSO negative control wells on each plate. Data plotted represents mean ± S.D. of six replicates (three technical replicates from two independent experiments).

### Dose-response (EC_50 ­­_and EC_90­_) for metabolites on *C. parvum* and host

To calculate metabolite EC_50_ values for *C. parvum* inhibition, metabolites were tested in a 7-point 1:2 serial dilution series starting at 200 μM (DCA, LCA and SDC), 4 mM (4HI and pyridoxal HCl), or 2 mM (indole and indole analogs). *C. parvum* growth assays were performed as described above, and relative parasite number was calculated as a ratio of the number of *C. parvum* in treated wells divided by the mean number of parasites in 1% DMSO negative control wells. To calculate EC­_50_ values for host cytotoxicity, the growth assay was performed with the following modifications: HCT-8 cells were plated at a lower concentration (5 x 10^4^ cells per well), cells were left uninfected, and the 7-point 1:2 serial dilution series started at 400 μM (DCA, LCA and SDC), 16 mM (4HI and pyridoxal HCl) or 8 mM (indole and indole analogs). Cells were fixed, permeabilized, stained with Hoechst and counted with the Cytation 3 as described above. Relative host toxicity was calculated as a ratio of the number of host nuclei in treated divided by the mean number of host nuclei in 1% DMSO negative control wells. EC_50_ and EC_90_ values for *C. parvum* and host cells were calculated using a nonlinear regression curve fit with six replicates per data point (three technical replicates from two independent experiments). Fold selectivity of a metabolite for parasite vs host was calculated as host EC_50_ divided by *C. parvum* EC_50._

### *C. parvum* growth assays with AhR agonists

AhR agonists VAF347 (Sigma) and FICZ (AAblocks, Inc.) were reconstituted at 10 mM in DMSO, while indole, 4-methylindole (4MeI) and kynurenic acid (Sigma) was reconstituted at 100 mM in DMSO. Compounds were diluted in culture medium to a starting concentration of 1 μM (VAF347 and FICZ) or 1 mM (indole, 4MeI, and kynurenic acid), then serially diluted 1:2 twice. *C. parvum* growth assays were performed as described above, and relative parasite numbers were calculated as a ratio of the number of *C. parvum* in treated wells divided by the mean number in 1% DMSO negative control wells. Data plotted represents mean ± S.D. of 9 replicates (three technical replicates from three independent experiments).

### Gene expression analysis of AhR target genes

HCT-8 cells were plated at 1 x 10^6^ cells per well in 12-well culture plates and grown for ∼24h until confluent. Medium in each well was then replaced with medium containing either 1% DMSO, VAF347 (250 nM), indole (1.5 mM) or 4-hydroxyindole (2.5 mM) and cultured for an additional 24h. Cells were then lysed in RLT buffer (QIAGEN) plus 1% ꞵ-mercaptoethanol and homogenized using a QIAshredder column (QIAGEN). RNA was extracted using the RNeasy Mini kit (QIAGEN), treated with RQ1 DNase (Promega) to remove DNA contamination, and converted to cDNA using the SuperScript VILO cDNA synthesis kit. Reverse transcription quantitative PCR (RT-qPCR) was run on a QuantStudio 3 real-time PCR system (Thermo Fisher Scientific) with TB Green Advantage qPCR premix (Takara Bio) and the following primers (5’ to 3’): human *CYP1A1* (forward, ACATGCTGACCCTGGGAAAG; reverse, GGTGTGGAGCCAATTCGGAT; PrimerBank ID 189339226c2), human *AHRR* (forward, CTTAATGGCTTTGCTCTGGTCG; reverse, TGCATTACATCCGTCTGATGGA^59^) and human *GAPDH* (forward, TGAGTACGTCGTGGAGTCCA; reverse, AGAGGGGGCAGAGATGATGA; this study). Relative gene expression was calculated with the ΔΔC_T_ method^60^ using human *GAPDH* as the reference gene and normalizing expression of each gene to its mean expression in the DMSO control samples. Data plotted represents mean ± S.D. of four technical replicates for all groups except for indole, which had 3 technical replicates due to RNA degradation in one of the samples.

### CRISPR/Cas9 KO of *AHR* in HCT-8 cells

A short guide RNA (5’-TCACCTACGCCAGTCGCAAG-3’) targeting the first exon of the human *AHR* gene (NM_001621) was cloned into BsmBI-digested Cas9 plasmid lentiCRISPR v2 (Addgene #52961). The resulting plasmid (pLentiCRISPRv2-sgAhR) was transfected along with support plasmids pMD2.g (Addgene #12259), pMDLg/pRRE (Addgene #12251), and pRSV-Rev (Addgene #12253) in equimolar concentrations into Lenti-X 293T cells (Takara Bio) using Lipofectamine 3000 reagent (Thermo Fisher Scientific). Lentivirus-containing supernatant was collected 72h post-transfection, passed through a 0.45 μM polyethersulfone (PES) filter, and diluted 1:2 in cell culture medium containing 10 μg/ml polybrene (Sigma) before adding to HCT-8 cells. Stable transgenic cells were selected for using puromycin (16 μg/ml) for 12 days starting 48 h after virus addition, with puromycin medium changes every 2 – 3 days. The transgenic cell population was serially diluted to isolate single cells, which were amplified and sequenced (Genewiz, Inc.) with primers flanking the targeted region (forward: 5’-GCACCATGAACAGCAGCAG-3’; reverse: 5’-TCCAAGTCCTCTGTCTCCCA-3’) to identify clonal lines with identical deleterious mutations in both alleles of *AHR*. Loss of AhR function was confirmed by treating AhR WT and KO cell lines with VAF347 (500 nM) or 1% DMSO for 24 h before harvesting RNA and analyzing *CYP1A1* gene expression as described above. Data plotted represents mean ± S.D. of three technical replicates per treatment group from one experiment.

*Cp* growth assay in HCT-8 WT vs AhR KO cell lines was performed as described above, treating with indole (1 mM), 4HI (1 mM), VAF347 (500 nM) or 0.5% DMSO. Data plotted represents mean ± S.D. of six replicates (three technical replicates from two independent experiments).

### Sliding window treatment with indoles

*C. parvum* growth assays were performed as described above with the following modifications: after allowing the HCT-8 cells to adhere to the bottom of the wells (4 h post-seeding), medium in “pre-treatment” wells was replaced with EC_90_ concentrations of indole (877 μM) or 7-cyanoindole (500 μM) in 0.87% DMSO medium. Medium in all other wells was replaced with 0.87% DMSO medium. After an additional 20h of culture, all wells were washed twice with medium and then infected with 1 x 10^5^ bleached oocysts. EC_90_ concentrations of indole or 7-cyanoindole in 0.87% DMSO medium were added to 0 – 4 h and 0 – 24 h treatment wells, with the remaining wells receiving 0.87% DMSO medium. All wells were washed twice with medium 4 hpi, and medium was replaced with EC_90_ concentrations of indole or 7-cyanoindole in 0.87% DMSO medium for the 4 – 24 h and 0 – 24 h treatment wells or 0.87% DMSO in medium for all other wells. All cells were fixed at 24 hpi and stained and imaged as described above. Relative parasite numbers were calculated as a ratio of the number of *C. parvum* in treated wells divided by the mean number in DMSO negative control wells. Data plotted represents mean ± S.D. of six replicates (three technical replicates from two independent experiments).

### Antibody and EdU labeling of *C. parvum*

HCT-8 cells were plated at 4.5 x 10^5^ cells per 12-mm-diameter glass coverslip (Thermo Fisher Scientific) in 24-well tissue culture plates and incubated until confluency (∼24 h). Monolayers were infected with ∼1 x 10^6^ excysted sporozoites, washed twice with DPBS at 4 hpi, then treated at EC_90_ concentrations of indole (880 μM) or 7-cyanoindole (500 μM) in 1% DMSO medium. For EdU pulse labeling, one set of two coverslips per treatment group was incubated with 10 μM EdU for 4 h, then fixed in 4% formaldehyde. EdU was then added to another set of two coverslips per treatment group for 4h before fixing, and this cycle was repeated until 20 hpi. All coverslips were permeabilized/blocked in 0.1% Triton-X + 1% BSA in DPBS for 20 min, then treated with the Click-iT Plus EdU 488 imaging kit (Thermo Fisher Scientific) for 30 min to label EdU. Parasites were labeled with mouse monoclonal antibody 1A5 and rabbit polyclonal antibody Pan Cp, followed by anti-mouse Alexa Fluor 568 (Thermo Fisher Scientific) and anti-rabbit Alexa Fluor 647 (Thermo Fisher Scientific) and Hoechst nuclear stain. Coverslips were mounted on glass slides using ProLong Glass antifade mountant (Thermo Fisher Scientific) and sealed with nail polish. The number of parasites at each life stage was counted from 10 fields using a 100X oil immersion objective on a Zeiss Axioskop Mot Plus fluorescence microscope. The sum of parasites at each life stage at each time point was divided by the total number of *C. parvum* for that time point. Ratios were averaged across three independent experiments.

For membrane labeling of *C. parvum* merozoites, HCT-8 cells were cultured on coverslips as described above and infected with 5 x 10^5^ *C. parvum* oocysts. Cells were washed twice 4 hpi with DPBS, and EC_90_ concentrations of indole (880 μM) or 7-cyanoindole (500 μM) in 1% DMSO medium were added. Monolayers were fixed and stained 22 hpi with mouse monoclonal antibody 1E12 and rabbit polyclonal antibody Pan Cp, followed by anti-mouse Alexa Fluor 488 (Thermo Fisher Scientific) and anti-rabbit Alexa Fluor 568 (Thermo Fisher Scientific) and Hoechst nuclear stain. Coverslips were mounted on glass slides using ProLong Glass antifade mountant (Thermo Fisher Scientific) and sealed with nail polish. Images were acquired on a Zeiss Axioskop Mot Plus fluorescence microscope with a 100X oil immersion objective using AxioVision software (Carl Zeiss, Inc.) and processed in ImageJ (https://fiji.sc/).

### Air-liquid interface cultures and washout assays

Air-liquid interface (ALI) cultures for long-term *C. parvum* growth were generated as previously described ^34,36^. Briefly, irradiated mouse 3T3 fibroblasts (i3T3) cells were plated on transwells (polyester membrane, 0.4-μm pore, 6.5 mm insert; Corning Costar) coated with 10% Matrigel (Corning) at a density of 8 x 10^4^ cells per transwell. Cells were cultured for 24h at 37 °C in DMEM high glucose medium supplemented with 10% fetal bovine serum, 100 U/ml penicillin, and 0.1 mg/ml streptomycin. Mouse intestinal epithelial cells (mIEC) spheroids were trypsinized and plated on the i3T3 feeder layer at 5 x 10^4^ mIECs per transwell and cultured in 50% L-WRN conditioned medium (CM) supplemented with 10 μM Y-27632 (ROCK inhibitor, RI), as defined previously^58,61^, with 200 μL and 400 μL medium in the top and bottom compartments of the transwell, respectively. Medium was changed every 2 to 3 days, and top medium was removed after 7 days to initiate the air-liquid interface.

To determine the indole and 7CNI EC_50_ and EC_90_ concentrations for *C. parvum* in ALI cultures, *C. parvum* oocysts (5 x 10^4^ per transwell) were added to the top of ALI cultures 3 days post top medium removal. After ∼4h of infection, the tops of the transwells were washed twice with DPBS to remove unexcysted oocysts, and the bottom medium was changed to medium containing indole or 7CNI in an 8 point serial dilution series. Approx. 48 hpi, cells were scraped from the transwell membranes, and DNA extraction was performed using the QIAamp DNA mini kit (QIAGEN). *C. parvum* and host genome equivalents were quantified on a QuantStudio 3 real-time PCR system (Applied Biosystems) with primers for their respective GAPDH genes, as previously described ^34^. EC_50_ and EC_90_ values were calculated using a nonlinear regression curve fit with seven replicates per data point (2-3 technical replicates from 3 independent experiments).

For indole and 7CNI washout experiments, *C. parvum* oocysts (5 x 10^4^ per transwell) were added to the top of ALI cultures 3 days post top medium removal. After ∼4h of infection, the tops of the transwells were washed twice with DPBS to remove unexcysted oocysts, and three transwells were scaped for DNA extraction (D0). The bottom medium in the remaining transwells was changed to medium containing 1% DMSO; indole at EC_50_ (577 μM), EC_90_ (1894 μM) or 2 x EC_90_ (3788 μM); or 7CNI at EC_50_ (379 μM), EC_90_ (688 μM) or 2 x EC_90_ (1376 μM). After ∼48 hours, the indoles were washed out by transferring transwells to clean wells with normal growth medium in the bottom compartments. A subset of transwells at each indole/7CNI concentration were scraped and collected for DNA at D2 (day of wash out), D4 (two days of recovery) and D6 (four days of recovery) post infection. *C. parvum* and host genome equivalents were quantified from DNA samples by qPCR as described above. Values from experimental samples were normalized to the mean of the DMSO control samples at each time point. Data plotted represents mean ± S.D. of six replicates (three technical replicates from two independent experiments).

### Transcriptomics of indole-treated HCT-8 cells

HCT-8 cells were plated at 9 x 10^5^ cells per well in 12-well culture plates and grown for ∼24h until confluent. Cells were infected with ∼4 x 10^6^ excysted *Cp* sporozoites per well and incubated for 4 h before washing 2X with DPBS to remove extracellular parasites. Cells were then treated with either indole (880 μM) or 1% DMSO in cell culture medium for an additional 4 h or 12 h before harvesting RNA. Cells were lysed in RLT buffer (QIAGEN) plus 1% ꞵ-mercaptoethanol (Sigma) and homogenized using a QIAshredder column (QIAGEN). RNA was extracted using the RNeasy Mini kit (QIAGEN) and treated with the DNA-*free* DNA Removal kit (Thermo Fisher Scientific). Total RNA from three technical replicates per treatment condition was submitted to the Genome Access Technology Center (Washington University School of Medicine) for library prep and sequencing. RNA integrity was determined using an Agilent Bioanalyzer (all samples had an RIN of 10) and library preparation was performed with 5 – 10 μg of total RNA per sample. Ribosomal RNA was removed by poly-A selection using Oligo-dT beads (mRNA Direct kit, Life Technologies). mRNA was then fragmented and reverse transcriptase buffer by heating to 94 °C for 8 min. mRNA was reverse transcribed using SuperScript III RT enzyme (Life Technologies) and random hexamers, per manufacturer’s instructions. A second strand reaction was performed to yield ds-cDNA, and then cDNA was blunt-ended before adding an A base to the 3’ ends and ligating Illumina sequencing adapters to both ends. Ligated fragments were amplified for 12 – 15 cycles using primers incorporating unique dual index tags, then sequenced on an Illumina NovaSeq 6000 to generate 150 bp, paired-end reads.

Demultiplexed fastq files were imported into Partek Flow (Partek, Inc.) and mapped to the Homo sapiens hg38 genome build (NCBI GenBank assembly ID GCA_000001405.15) using the STAR 2.7.8a aligner with default parameters^62^. The number of reads mapping to each gene was then quantified based on the human Ensembl Transcripts release 100. At this point, we removed one of the replicates treated with DMSO for 4 h from further analysis due to most of its reads mapping to intergenic regions (indicative of DNA contamination). For the remaining 11 samples, gene expression values were normalized across samples by dividing the number of reads per gene by the total number of reads per sample to obtain counts per million (CPM) values. A pair-wise comparison of indole-treated (6 total) and DMSO-treated samples (5 total) was performed using the Partek gene-specific analysis (GSA) algorithm using default parameters. Volcano plots were generated in Partek Flow. Genes were considered significantly differentially expressed if the FDR-corrected *P* value was less than 0.05 and the absolute fold change was greater than 2. Hierarchical clustering analysis of significant genes was performed in Partek Flow using a Euclidean point distance matrix and average linkage for the cluster distance metric. Gene Ontology (GO) pathway analyses were performed in Enrichr (http://maayanlab.cloud/Enrichr/)^63-65^ with the GO Biological Process 2021 database.

### *C. parvum* growth assay with amino acid supplementation

HCT-8 cell culture medium was supplemented with 1 mM L-tryptophan (Sigma) with or without 1 mM L-phenylalanine (Sigma), and then filter sterilized. Indole was tested in a 5-point 1:2 serial dilution starting at 2 mM in all three media. *C. parvum* growth assays were performed as described above, and relative parasite number was calculated as a ratio of the number of *C. parvum* in each well divided by the mean number of parasites in the 1% DMSO, non-supplemented medium control wells. Indole EC_50_ values on *Cp* in each medium type were calculated using a nonlinear regression curve fit (with six replicates per data point (three technical replicates from two independent experiments).

### Metabolic assays using Seahorse

For all metabolic assays, mitochondrial oxygen consumption rates (OCR) and extracellular acidification rates (ECAR) were measured using a Seahorse XF96 Analyzer. HCT-8 cells or HCT-8 AhR KO cells were plated at 2 x 10^4^ cells per well in a 96-well Seahorse XF96 cell culture microplate (Agilent) and grown for ∼24h. Medium was replaced with fresh medium containing either 1% DMSO or 1:2 serial dilutions of indole (starting at 2 mM) or 7CNI (starting at 1 mM) and cultured for an additional 18h (cell confluency ∼90%). A Seahorse XFe96 extracellular flux sensor cartridge (Agilent) was hydrated overnight in distilled water in a non-CO_2_, 37 °C incubator. The water was then replaced with XF Calibrant (Agilent), and the sensor cartridge was incubated for an additional ∼1h in a non-CO_2_, 37 °C incubator. Assay medium was prepared by adding 1 mM pyruvate (Agilent), 2 mM glutamine (Agilent) and 10 mM glucose (Agilent) to Seahorse XF DMEM medium, pH 7.4 (Agilent).

For assays run with the Seahorse XF Cell Energy Phenotype kit (Agilent), cells were washed once with assay medium before incubating in assay medium in a non-CO_2_, 37 °C incubator for 1h. OCR and ECAR levels were measured for five cycles of mixing (2 mins) and measuring (3 mins) before and after supplementation via injection ports with a mix of FCCP (1 μM) and oligomycin (1 μM).

For assays run with the Seahorse XF Real-time ATP Rate Assay kit (Agilent), cell growth medium was replaced with assay medium and incubated in a non-CO_2_, 37 °C incubator for 1h. Assay medium was then replaced with fresh assay medium prior to running the assay. OCR and ECAR levels were measured for four cycles of mixing (2 mins) and measuring (3 mins) before and after sequential supplementation via injection ports with oligomycin (1.5 μM), followed by a mix of rotenone (0.5 μM) and antimycin A (0.5 μM).

For assays run with the Seahorse XF Cell Mito Stress Test Kit (Agilent), cells were washed once with assay medium before incubating in assay medium in a non-CO_2_, 37 °C incubator for 1h. OCR and ECAR levels were measured for four cycles of mixing (2 mins) and measuring (3 mins) before and after sequential supplementation via injection ports with oligomycin (1.5 μM), followed by FCCP (1 μM), and then a mix of rotenone (0.5 μM) and antimycin A (0.5 μM).

Results were initially analyzed in Wave v2.6.1 (Agilent) before importing into Graphpad Prism 9. All assays were performed twice with six technical replicates per sample per plate. Wells were removed from analysis if the OCR or ECAR values were negative or identified as an outlier by a ROUT test with Q = 1% performed in GraphPad Prism 9. Data plotted represents mean ± S.D. of 10 – 12 replicates (5 – 6 technical replicates from two independent experiments).

### *C. parvum* growth assay with mitochondrial inhibitors

Rotenone (Sigma), antimycin A (Sigma), oligomycin (Sigma) and carbonyl cyanide m-chlorophenyl hydrazone (CCCP, Sigma) were reconstituted at 20 mM in DMSO. Compounds were diluted in DMSO to 4 mM each with rotenone and antimycin-A mixed together, each at 4 mM. Compounds were serially diluted 1:2 7X in DMSO, then diluted in cell culture medium to 1% DMSO with the highest concentration of each compound starting at 40 μM. *C. parvum* growth assays were performed as described above, and relative parasite numbers were calculated as a ratio of the number of *C. parvum* in treated wells divided by the mean number in 1% DMSO negative control wells. Data plotted represents mean ± S.D. of 4 replicates (two technical replicates from two independent experiments). Data for CCCP at 40 μM and 20 μM were not included due to observed host toxicity at those concentrations.

### Mitotracker staining of indole treated *C. parvum*

HCT-8 cells were cultured on coverslips and infected with 5 x 10^5^ *C. parvum* oocysts. Cells were washed twice with DPBS and medium was supplemented with 2 x EC_90_ concentrations of indole (1.76 mM), 7-cyanoindole (1 mM) or 10 μM carbonyl cyanide m-chlorophenyl hydrazone (CCCP, Sigma) in 1% DMSO media. Infected cells were treated for 2 h (22 – 24 hpi) or continuously for 24 hpi vs. 1% DMSO control. MitoTracker Red CMXRos (50 nM final concentration) was added to the culture 45 mins before fixation. Monolayers were fixed and stained at 24 hpi with mouse monoclonal antibody 1E12 followed by anti-mouse Alexa Fluor 488 (Thermo Fisher Scientific) and Hoechst nuclear stain. Coverslips were mounted on glass slides using ProLong Glass antifade mountant (Thermo Fisher Scientific) and sealed with nail polish. Images were acquired on a Zeiss Axioskop Mot Plus fluorescence microscope with a 100X oil immersion objective using AxioVision software (Carl Zeiss, Inc.) and the relative fluorescence intensity was processed in ImageJ (https://fiji.sc/)^66^. Example parasites were chosen from four replicates (two technical replicates from two independent experiments).

### Metabolite treatment of *C. parvum*-infected GKO mice

In the first set of *in vivo* metabolite experiments, indole and 7-cyanoindole solutions were prepared at 5 mg/ml in vehicle (10% DMSO and Millipore-filtered water), aliquoted and stored at −20 °C. A total of 9 GKO mice between 2 – 4 months old were split into 3 treatment groups: vehicle BID (1 male, 2 females), 50 mg/kg BID indole (1 male, 2 females) and 50 mg/kg BID 7CNI (2 males, 1 female). Within each treatment group, mice were co-housed with siblings but separated by sex. On the first day of the experiment (0 dpi), all mice were infected by oral gavage with 2 x 10^4^ *Cp* oocysts mixed in 200 µl of their respective treatment solution. Mice received an additional 200 µl gavage of their treatment solution approx. every 12 h for a total of 7 days. Mice were weighed every 24 – 48 hpi and survival was recorded. A single fecal pellet was collected per mouse 3, 5, 7, and 9 dpi. Fecal pellets were stored at −80 °C until further processing. The experiment was terminated 10 dpi.

In the second set of *in vivo* metabolite experiments, indole and 7-cyanoindole solutions were prepared at 10 mg/ml in vehicle (10% DMSO and Millipore-filtered water), aliquoted and stored at −20 °C. A total of 12 GKO mice between 2 – 3 months old were split into the same 3 treatment groups as previously (vehicle BID, 50 mg/kg BID indole, and 50 mg/kg BID 7CNI) with 2 males and 2 females per group. Within each treatment group, mice were co-housed with siblings but separated by sex. On the first day of the experiment (0 dpi), all mice were infected by oral gavage with 2 x 10^4^ *Cp* oocysts in 100 uL PBS. Approx. 30 mins later, all mice received 100 µl of their respective treatment solution via oral gavage. Mice received an additional 100 µl gavage of their treatment solution approx. every 12 h for a total of 7 days. Mice were weighed every 24 – 48 hpi and survival was recorded. A single fecal pellet was collected per mouse 3, 5, 7, and 9 dpi. Fecal pellets were stored at −80 °C until further processing. The experiment was terminated at 30 dpi.

DNA was extracted from fecal pellets using the QIAamp Powerfecal Pro DNA kit (QIAGEN) following the manufacturer’s protocol. The number of *Cp* oocysts per fecal pellet was calculated using qPCR for the *Cp* GAPDH gene as described above and then divided by the weight of the pellet in mg to obtain the number of *Cp* per mg feces.

### *B. theta* reconstitution of *C. parvum*-infected GKO mice

GKO mice between 8-16 weeks old were separated by sex and split into 3 treatment groups: vehicle (2 males, 2 females), WT *B. theta* (2 males, 2 females) and *ΔtnaA B. theta* (2 males, 2 females). Within each treatment group, mice were co-housed with siblings but separated by sex. All mice were given water (Abx water) containing 1 g/L ampicillin, 1 g/L neomycin and 0.5 g/L vancomycin (Sigma) for 3 days to deplete gut microbiota. Following Abx treatment (−3 dpi), groups of mice were inoculated with WT *B. theta* or with *ΔtnaA B. theta* (1 x 10^8^ CFU) while control mice were orally administered vehicle (10% glycerol in Millipore-filtered water). Mice received three additional treatments at −2, 3, 5 dpi to reconstitute gut microbiota. On the first day of the experiment (0 dpi), all mice were infected by oral gavage with 2 x 10^4^ *Cp* oocysts in 100 μl PBS. Fecal pellets were collected at 3, 5, 7, and 9 dpi for measuring *Cp* oocysts in feces, and at −6, −3, 0, 4, 8 dpi for analyzing gut microbiota. Fecal pellets were stored at −80 °C until further processing. The experiment was terminated at 16 dpi and mice were humanely sacrificed. DNA was extracted from fecal pellets using the QIAamp Powerfecal Pro DNA kit (QIAGEN) following the manufacturer’s protocol. The number of *Cp* oocysts per fecal pellet was calculated using qPCR for the Cp GAPDH gene as described above and then divided by the weight of the pellet in mg to obtain the number of *Cp* per mg feces.

### 16S rRNA analyses

For quantification of bacterial levels, the 16S rRNA gene was amplified by qPCR and quantified using SYBR Green. Primers for PCR consisted of 515 F and 805 R to detect the V4 hypervariable region of the 16S rRNA gene: (forward, 5’-GTGCCAGCMGCCGCGGTAA-3’; reverse, 5’-GACTACCAGGGTATCTAATCC-3’). For 16S rRNA sequencing, PCR amplification of the V4 region was performed as described previously ^67^. Briefly, each sample was amplified in triplicate with Golay-barcoded primers specific for the V4 region (F515/R805) and confirmed by gel electrophoresis. Platinum High Fidelity Taq (Invitrogen) was used for PCR. Amplicons were pooled and purified with 0.6x Agencourt Ampure XP beads (Beckman-Coulter) according to the manufacturer’s instructions. The final pooled samples, along with aliquots of the three sequencing primers, were sequenced using a 2 x 250 bp Illumina protocol by the DNA Sequential Innovation Lab (Washington University School of Medicine). Read quality control and the resolution of amplicon sequence variants (ASVs) were performed with the dada2 R package ^68^. ASVs that were not assigned to the kingdom Bacteria were filtered out. The remaining reads were assigned taxonomy using the Ribosomal Database Project (RDP trainset 16/release 11.5) 16S rRNA gene sequence database ^69^. Ecological analyses, such as alpha-diversity (richness) and beta-diversity analyses (weighted UniFrac distances), were performed using PhyloSeq and additional R packages ^70^.

### Statistical Analyses

All statistical analyses were performed in GraphPad Prism 9. A one-way ANOVA (single variable) or two-way ANOVA (multiple variables) followed by a Dunnett’s multiple comparisons test was used to compare each experimental group to a control group (defined in each figure legend). EC_50_ and EC_90_ values were calculated using a nonlinear regression curve fit (log inhibitor vs normalized response – variable slope), with the number of replicates specified in the methods above. For mouse experiments, each treatment group was compared to control group on individual days using a two-tailed Mann-Whitney U test. Comparisons of body weight between treatment groups was analyzed using a mixed-effects model with a Geisser-Greenhouse correction for matched values, followed by a Dunnett’s test for multiple comparisons.

### Data Availability

Raw RNA-seq reads and analyzed data generated in this study have been deposited in the Gene Expression Omnibus (GEO) database under accession number GSE185652. Sequence reads for the 16S RNA analyses have been deposited in the European Nucleotide Archive and are available under accession PRJEB58007.

## ACKNOWLEDGMENTS

We thank William Witola (University of Illinois at Urbana-Champaign) for providing the *C. parvum* oocysts and Vanessa Sperandio for providing the *B. theta* strains used here and Lynn Foster for technical assistance. Support was provided in part by grants from the NIH (AI 145496 to L.D.S. and DK109081 to K.L.V.). We thank the Genome Technology Access Center at the McDonnell Genome Institute at Washington University School of Medicine for help with genomic analysis. The Center is partially supported by NCI Cancer Center Support Grant #P30 CA91842 to the Siteman Cancer Center and by ICTS/CTSA Grant# UL1TR002345 from the National Center for Research Resources (NCRR), a component of the National Institutes of Health (NIH), and NIH Roadmap for Medical Research. This publication is solely the responsibility of the authors and does not necessarily represent the official view of NCRR or NIH.

