## Supplementary material for "Microbiota produced indole metabolites disrupt host cell mitochondrial energy production and inhibit *Cryptosporidium parvum* growth"

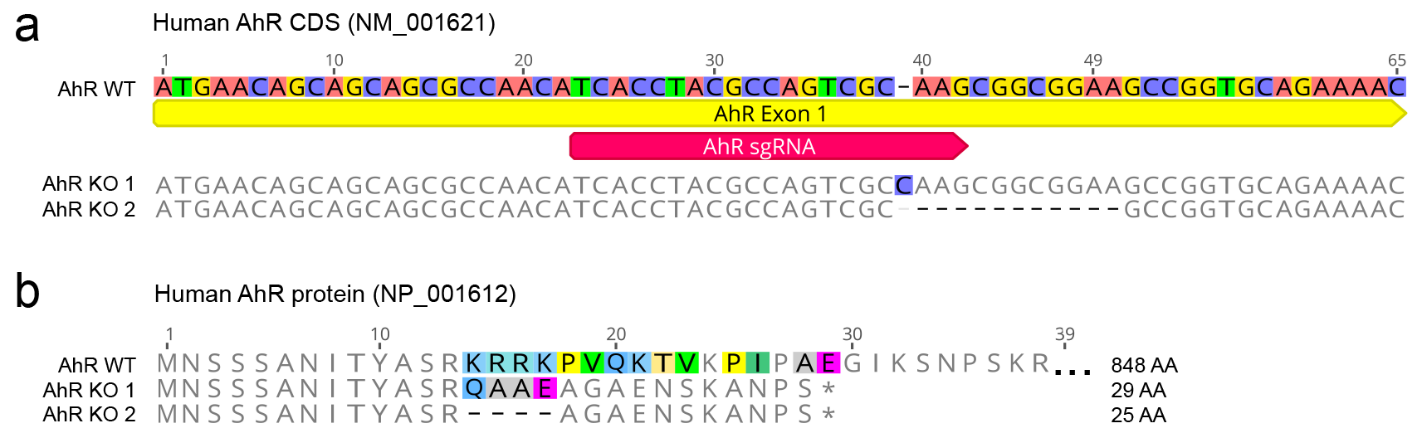

**Supplementary Fig. 1: Mutations in the AhR gene introduced by CRISPR/Cas9 cause frameshift mutations resulting in truncated proteins.** a) Nucleotide alignment of exon 1 for the WT human AhR gene (NM_001621) with sequences from two mutated HCT-8 cell lines, one with a single bp insertion highlighted in blue (AhR KO 1) and the other with an 11 bp deletion (AhR KO 2). b) Amino acid (AA) alignment of the first 38 AA of the WT human AhR protein (NP_001612) with the predicted truncated protein sequences for the AhR KO lines. Graphics were created in Geneious v.2021.1.

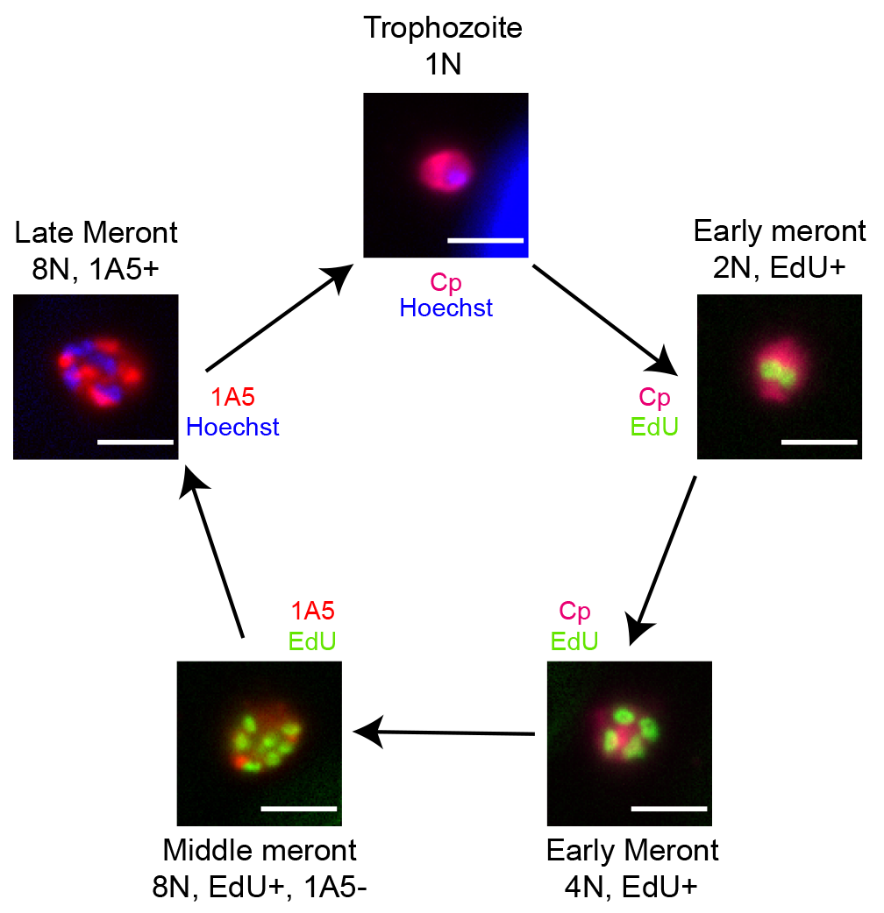

**Supplementary Fig. 2**: **Staging *C. parvum* cell cycle progression.** Asexual stages were defined based on the number of nuclei (N), EdU incorporation and/or labeling of mature merozoites with monoclonal antibody 1A5, as follows: trophozoites had 1 nucleus, early meronts had either 2 or 4 EdU+ nuclei, middle meronts had 8 EdU+ nuclei but no 1A5 labeling, and mature meronts had 8 nuclei with 1A5 labeling.

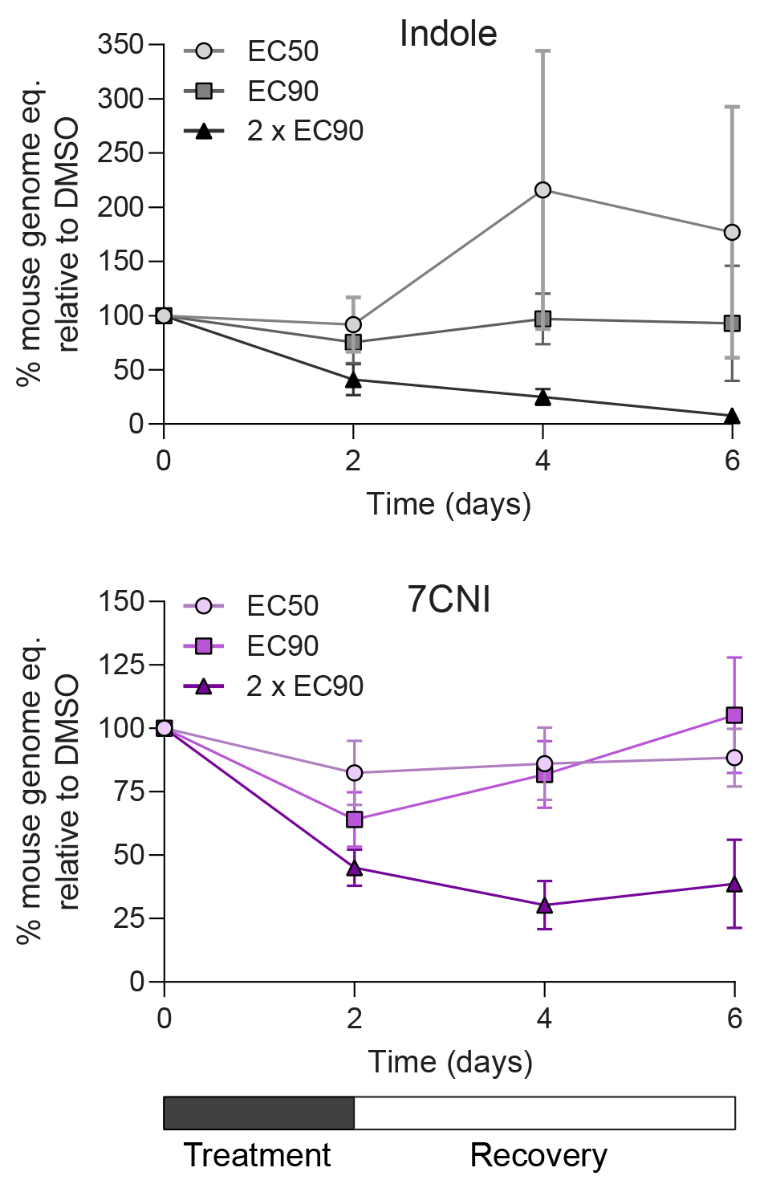

**Supplementary Fig. 3: High concentrations of indole and 7-cyanoindole are toxic to host cells in an air-liquid interface (ALI) transwell culture system**. Washout experiments in *Cp*-infected ALI cultures treated with 1% DMSO or indole at EC_50_ (577 μM), EC_90_ (1894 μM) or 2 x EC_90_ (3788 μM); or 7CNI at EC_50_ (379 μM), EC_90_ (688 μM) or 2 x EC_90_ (1376 μM) for 48 h before washout. Mouse genome eq. were normalized to the DMSO control at each time point. Data plotted represents mean ± s.d. of six replicates (three technical replicates from two independent experiments).

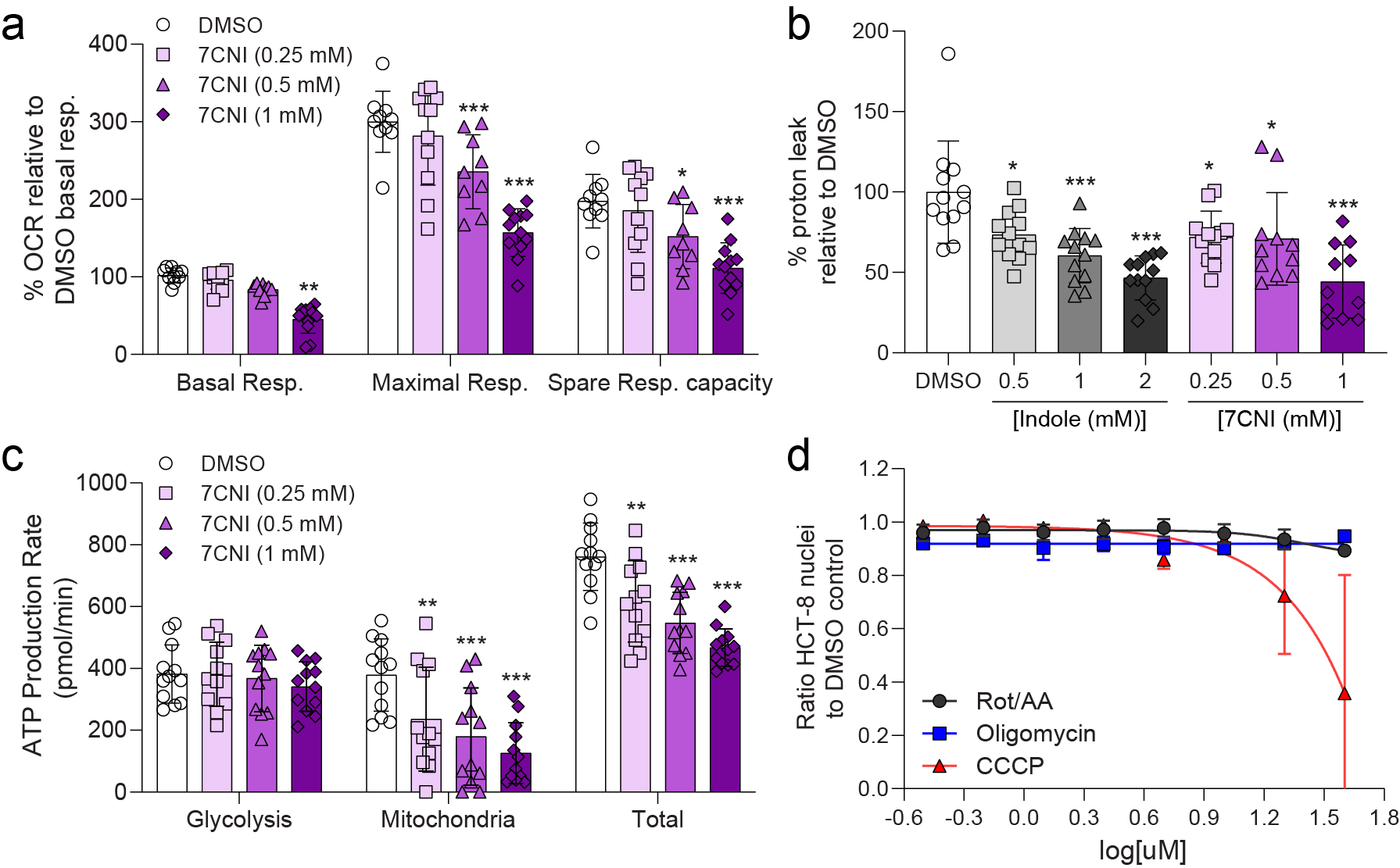

**Supplementary Fig. 4: Indoles impair host mitochondrial ATP production but not through proton leak.** a) Metabolic analysis using the Seahorse XF Cell Mito Stress Test kit on HCT-8 cells treated for 18 h with 1% DMSO or 7-cyanoindole (7CNI, 0.25 mM, 0.5 mM or 1 mM). Data calculated as a percentage of the oxygen consumption rate (OCR) for each well relative to the mean basal OCR of DMSO control cells for that experiment. Spare respiratory capacity = maximal respiratory rate – basal respiratory rate for each well. Data plotted represents mean ± s.d. of 12 replicates (six technical replicates from two independent experiments). Differences between *%* OCR for each indole concentration vs the DMSO control for each measurement were analyzed with a two-way ANOVA followed by a Dunnett’s test for multiple comparisons. **P* < 0.05, ****P* < 0.001. b) Metabolic analysis using the Seahorse XF Cell Mito Stress Test kit on HCT-8 cells treated for 18 h with 1% DMSO, indole (0.5 mM, 1 mM or 2 mM) or 7CNI (0.25 mM, 0.5 mM or 1 mM). Data calculated as a percentage of the proton leak for each well relative to the mean proton leak of DMSO control cells for that experiment. Data plotted represents mean ± s.d. of 12 replicates (six technical replicates from two independent experiments). Differences between *%* proton leak for each indole/7CNI concentration vs the DMSO control were analyzed with a one-way ANOVA followed by a Dunnett’s test for multiple comparisons. **P* < 0.05, ****P* < 0.001. c) Metabolic analysis using the Seahorse XF Real-time ATP Rate assay on HCT-8 cells treated for 18 h with 1% DMSO or 7CNI (0.25 mM, 0.5 mM or 1 mM). Data plotted represents mean ± s.d. of ATP production rate (pmol/min) produced by glycolysis, the mitochondria, or total ATP (glycolysis + mitochondrial ATP rates) for 12 replicates (six technical replicates from two independent experiments). For each source of ATP, differences between ATP production rate for each 7CNI concentration vs the DMSO control were analyzed with a two-way ANOVA followed by a Dunnett’s test for multiple comparisons. ***P* < 0.01, ****P* < 0.001. c) d) Ratio of host nuclei relative to DMSO control in HCT-8 cells after 24 h treatment with serial dilutions of mitochondrial Complex I and III inhibitors rotenone and antimycin A, respectively (Rot/AA); ATP synthase inhibitor oligomycin; or proton gradient uncoupler carbonyl cyanide m-chlorophenyl hydrazone (CCCP). Inhibition curves were calculated for each compound using a nonlinear regression curve fit with six replicates (three technical replicates from two independent experiments) per concentration.

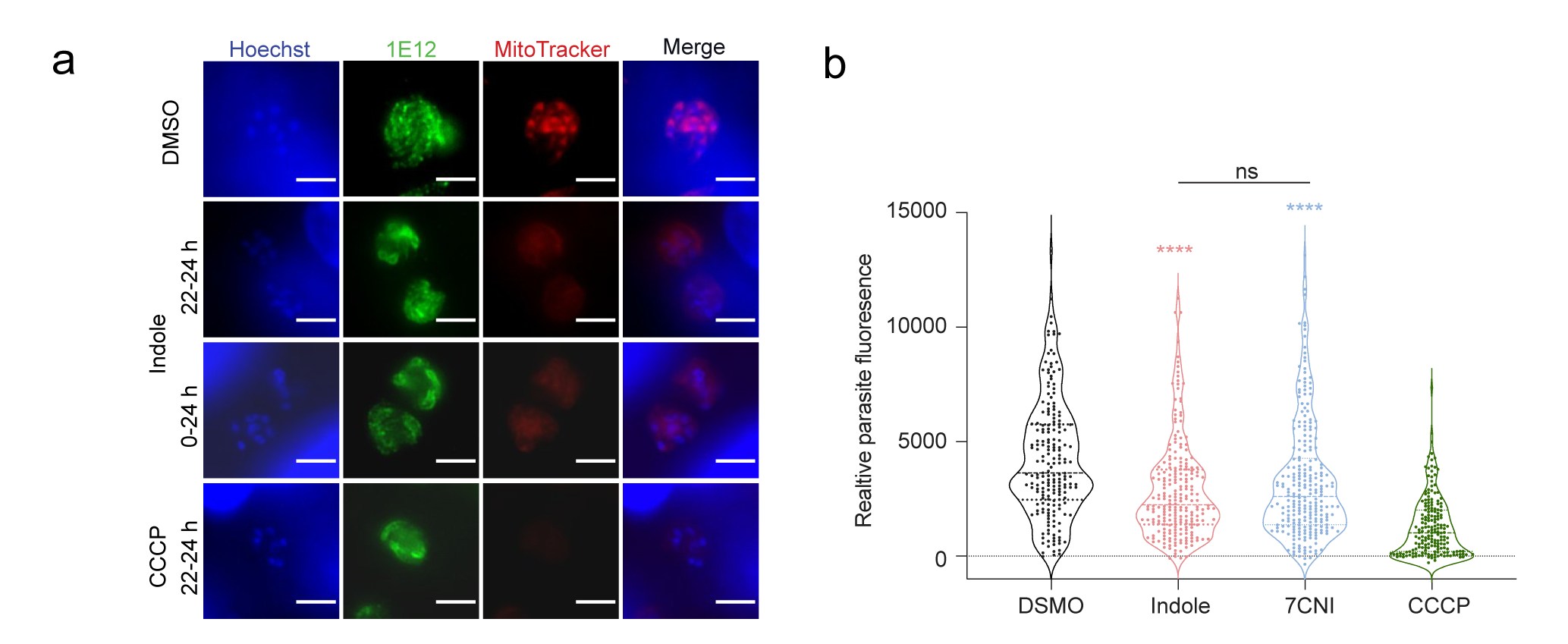

**Supplementary Fig. 5** **Treatment with indole affects the membrane polarity of the mitosome.** a) Immunofluorescence images of *Cp* in HCT-8 cultures treated with 1% DMSO or 2 x EC_90_ concentrations of indole (1.76 mM) or 10 μM CCCP for the indicated hpi. Parasites were labeled with membrane marker 1E12 (green), MitoTracker Red CMXRos (red), and nuclei were stained with Hoechst (blue). Scale bar = 3 μm. b) Distribution of MitoTracker intensity for indole treated parasites. *Cp* in HCT-8 cultures treated with 1% DMSO or 2 x EC_90_ concentrations of indole (1.76 mM), 7-cyanoindole (7CNI) (1 mM) or 10 μM CCCP for 2 hours starting 22 hpi. Parasite fluorescence intensity was measured based on MitoTracker staining collected from at least 180 parasites from two independent experiments. Statistical analyses comparing each treatment group to control were performed using two-tailed Mann-Whitney U tests. *****P* < 0.0001. ns = not significant. (DMSO compared to indole shown in red asterisk, DMSO compared to 7CNI shown in blue asterisk.)

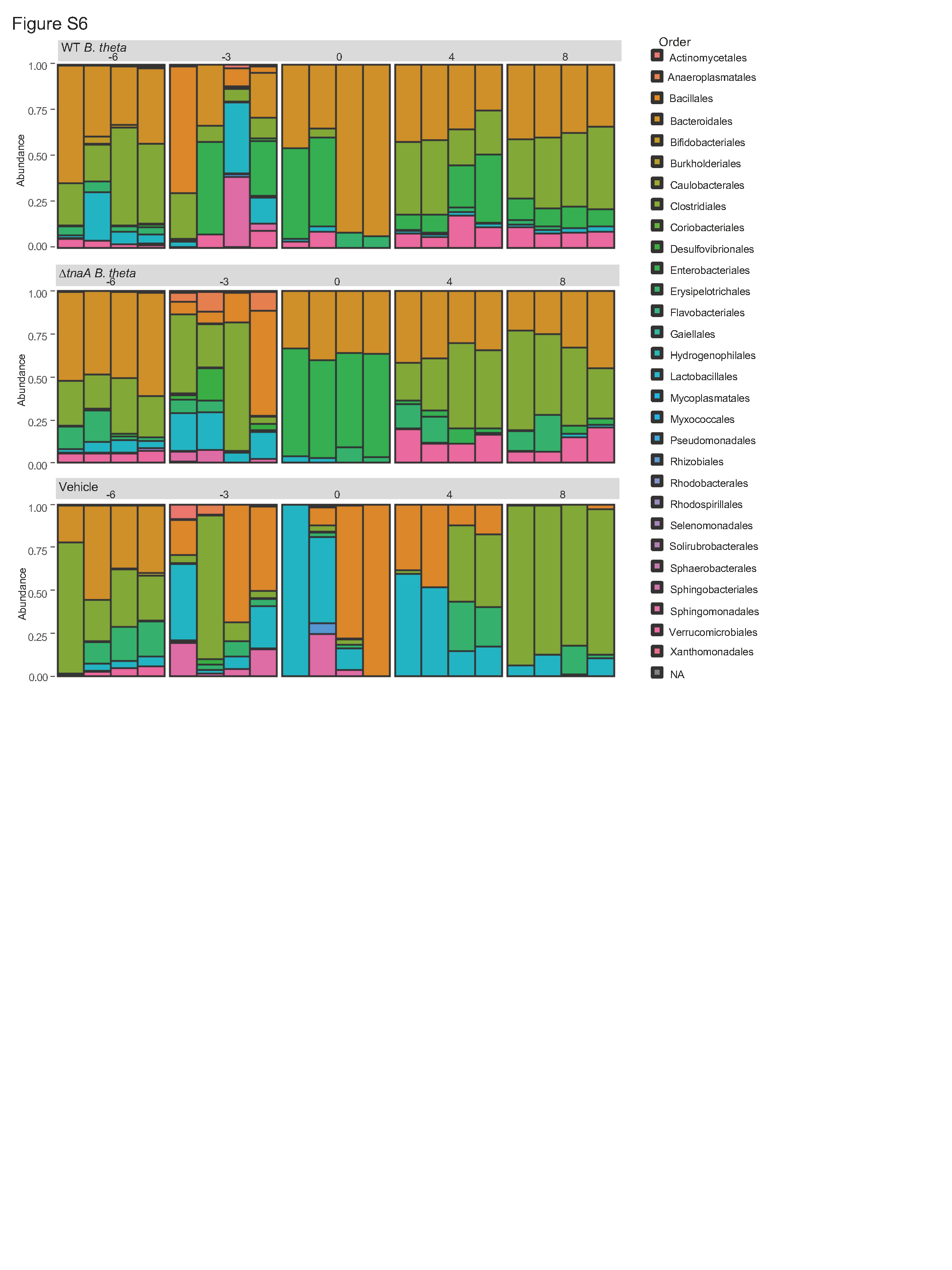

**Supplementary Fig. 6: 16S profiling of commensal microbiota in fecal pellets**. Order level composition for each group of mice at the indicated days post infection. Data represent 4 mice per treatment group sampled over time. Prior to antibiotic treatment, Bacteroidales were the major component of the microbiota. Following antibiotic treatment and reconstitution, *B. theta* replaced the exiting taxa and consequently Bacteroidales was the most common order in mice given wild type (WT) *B. theta* and the *∆tnaA* mutant bacteria. In contrast, in mice given the vehicle control the major order was Bacillales.

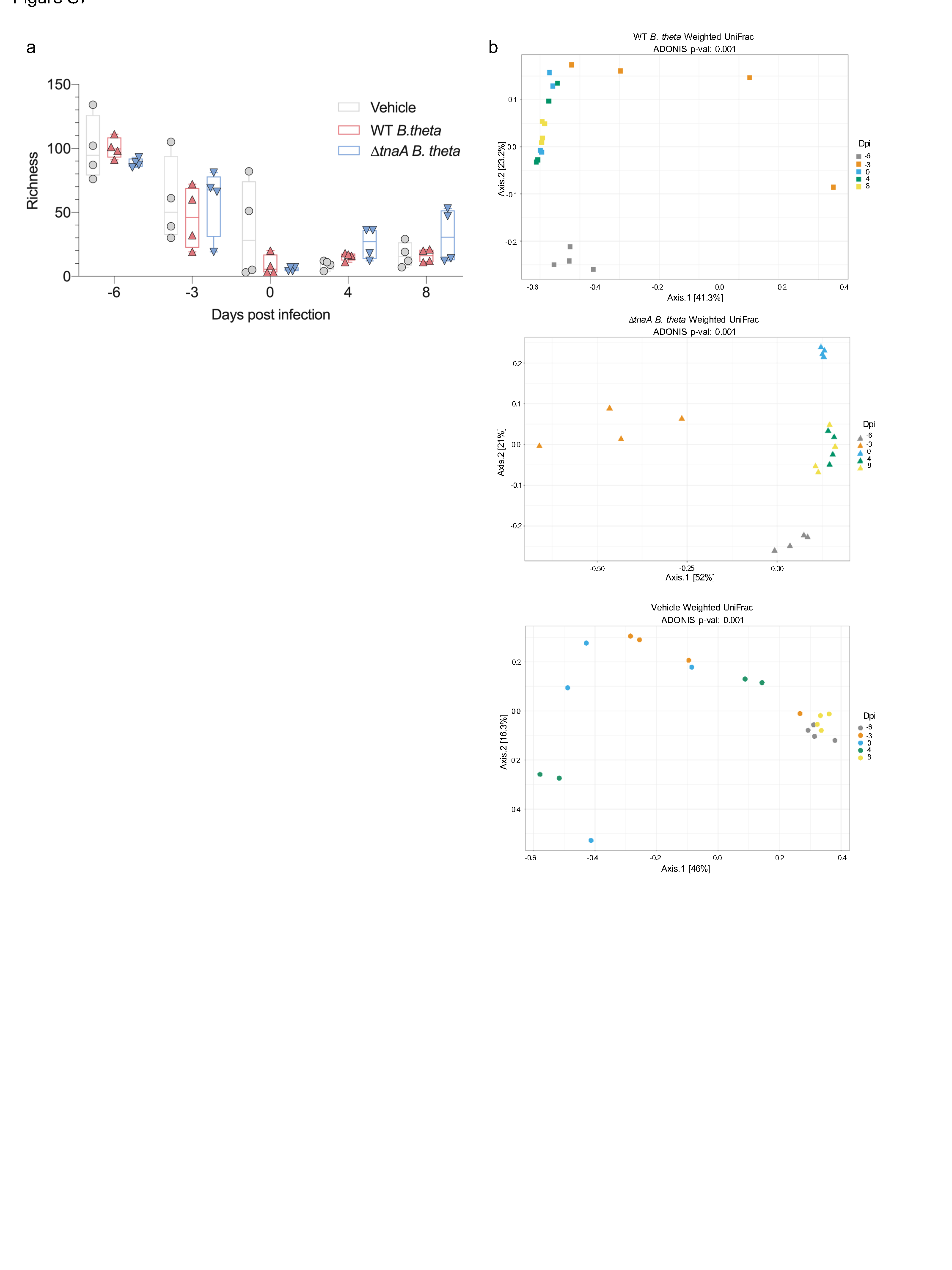

**Supplementary Fig. 7: 16s rRNA sequencing analysis in fecal pellets.** a) Alpha diversity for each group of mice at the indicated dpi. Principal components analysis was using richness diversity. Statistical analyses comparing vehicle to WT *B. theta* group or *∆tnaA* *B. theta* group on individual days were analyzed with a two-way ANOVA followed by a Dunnett’s test for multiple comparisons. All of the comparison showed no significant difference. b) Beta diversity plot for each group of mice at the indicated dpi. Principal components analysis was performed using uniFrac distance. Differences in beta diversity of each group were analyzed using Permutational Multivariate Analysis of Variance (ADONIS). All data plotted represents 4 mice per treatment group sampled over time.
